# Latent Transforming Growth Factor Binding Protein-2 promotes microfibrillar TGF-β bioavailability to drive pathologic fibrosis in remote regions of infarcted hearts

**DOI:** 10.64898/2026.10.28.762676

**Authors:** Fahad Ehsan, Michelle Di Paola, Michael B. Dewar, Hangjun Zhang, Uros Kuzmanov, Yu-Qing Zhou, Cristine J. Reitz, Haisam Shah, Tao Wang, Jingzhi Liu, Aliya Izumi, Rundong Hu, Joanna Hendy, Fereshteh S. Younesi, Mark Friedberg, Filio Billia, Tomoyuki Nakamura, Boris Hinz, Robert P. Mecham, Anthony O. Gramolini, Scott P. Heximer

## Abstract

**Background:** Myocardial infarction (MI) initiates ventricular remodeling that extends beyond the infarct into remote, viable myocardium, where diffuse extracellular matrix (ECM) remodeling and stiffening contribute to progressive dysfunction and heart failure. Latent transforming growth factor binding protein 2 (LTBP2) is an injury-induced, microfibril-associated ECM glycoprotein that, uniquely among the LTBP family, does not covalently bind latent TGF-β. We asked whether LTBP2 regulates remote remodeling after MI, and how a protein lacking latent TGF-β–binding capacity could influence TGF-β–dependent fibrosis.

**Methods:** Wild-type and *Ltbp2*-null mice underwent permanent coronary ligation or sham surgery and were followed for 28 days by echocardiography and speckle-tracking strain analysis. Remote myocardium was assessed by histology, atomic force microscopy, and quantitative proteomics. Cellular mechanism was interrogated in primary cardiac fibroblasts (CFs) using a TGF-β reporter assay and quantitative imaging of LTBP1 and LTBP2 matrix incorporation. Sufficiency was tested with a minimal fibrillin-1–binding module of LTBP2 delivered as purified protein or by lentiviral expression.

**Results:** LTBP2 was induced in injury-activated CFs and localized to infarct and remote myocardium by 7 days post-MI. *Ltbp2* deletion improved 28-day survival (P=0.0272) without altering infarct size or early post-infarct function. By 28 days, *Ltbp2*-null hearts showed improved ejection fraction and faster diastolic strain rates, with protection concentrated in segments remote from the infarct. Remote myocardium contained less interstitial collagen and was significantly less stiff, while cardiomyocyte hypertrophy was unaffected. Proteomic profiling showed enrichment of ECM organization and TGF-β–responsive programs in wild-type hearts that was attenuated in knockouts. In fibroblasts, loss of LTBP2 increased LTBP1 matrix incorporation and shifted TGF-β from the soluble to the matrix-bound compartment; conversely, the minimal LTBP2 module bound fibrillin-1 directly and mobilized latent TGF-β from established matrix within 20 minutes.

**Conclusions:** LTBP2 drives maladaptive remodeling of remote myocardium after MI by limiting matrix sequestration of latent TGF-β and expanding the activatable pool, resolving how an LTBP that does not bind latent TGF-β amplifies TGF-β–dependent fibrosis. LTBP2 and its fibrillin-1 interaction surface may be tractable targets.

**Graphical Abstract:** **LTBP2 remodels microfibrillar composition to enhance TGF-β bioavailability and profibrotic signaling.** Under homeostatic conditions (left), latent TGF-β is sequestered within the extracellular matrix (ECM) as part of the large latent complex (LLC), in which TGF-β is covalently associated with LTBP1 or LTBP4 and tethered to fibrillin-1 microfibrils. This matrix sequestration maintains a balanced distribution between bound and soluble pools of latent TGF-β, thereby limiting ligand activation and downstream signaling. During fibroblast activation (right), LTBP2 expression is upregulated and incorporated into newly assembling fibrillin-1 microfibrillar networks. Although LTBP2 does not covalently bind latent TGF-β, it competes with TGF-β–binding LTBP isoforms for fibrillin-1 binding sites, reducing LLC deposition within the matrix. This competitive remodeling shifts the equilibrium from matrix sequestration toward an expanded soluble LLC pool. Soluble latent complexes are more readily activated through integrin-mediated mechanical stress or proteolytic processing, increasing local active TGF-β levels. This model provides a mechanistic framework whereby LTBP2 indirectly amplifies TGF-β signaling by altering microfibrillar composition and competitive binding dynamics within the ECM, promoting enhanced TGF-β bioavailability and sustained profibrotic remodeling in the remote myocardium.

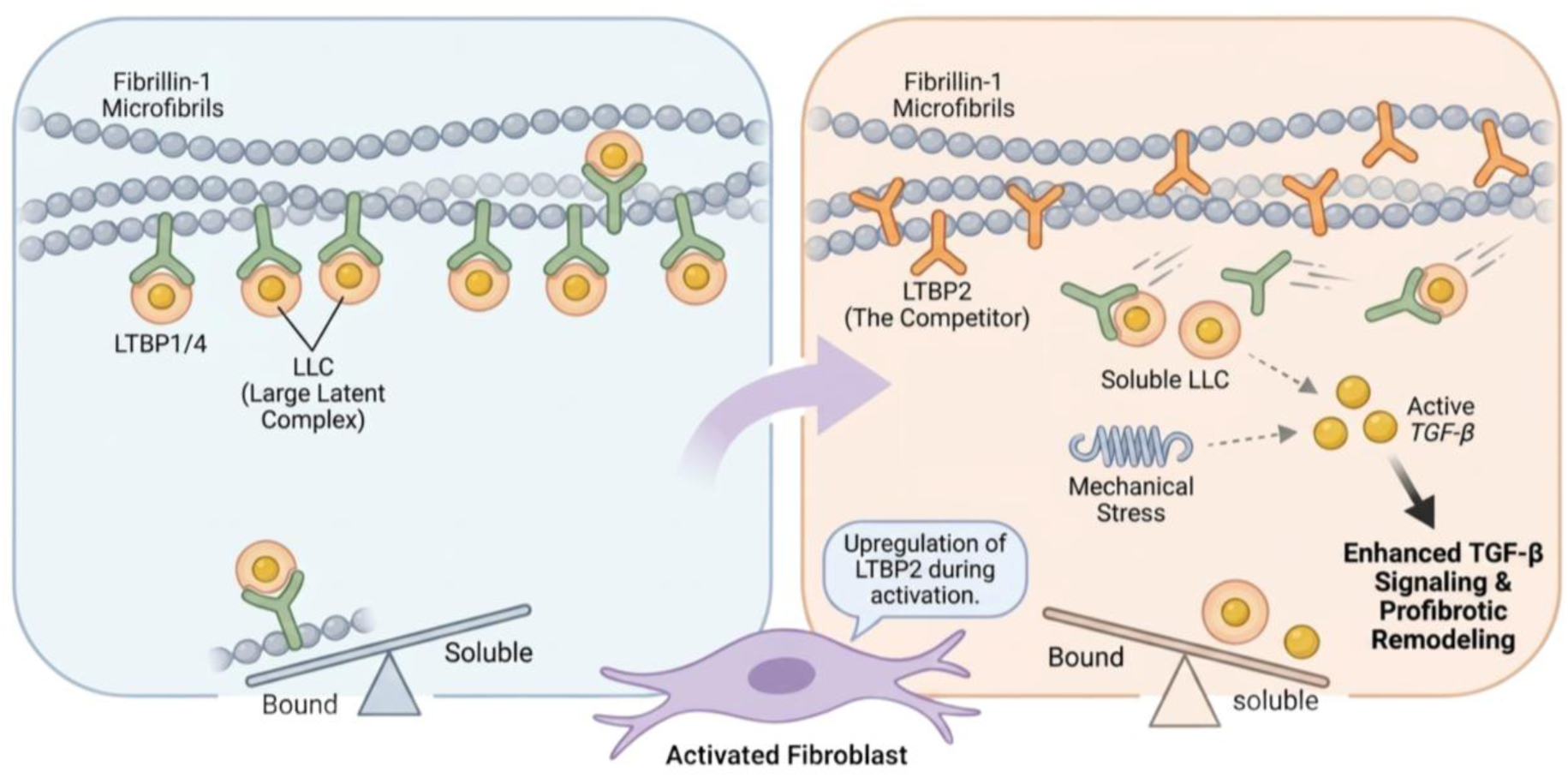

## Introduction

Myocardial infarction (MI) triggers a coordinated wound-healing response that remodels the left ventricle well beyond the ischemic scar. Although necrosis, inflammation, and replacement fibrosis within the infarct zone are well characterized, an increasingly important determinant of long-term outcome is remodeling of the remote, viable myocardium (1–3). Remote remodeling can include cardiomyocyte hypertrophy, diffuse interstitial fibrosis, microvascular dysfunction, mitochondrial impairment, and persistent inflammatory and neurohormonal activation, all of which contribute to progressive ventricular dilation and contractile dysfunction (4–5). Clinical and pre-clinical studies show that remote myocardial abnormalities correlate strongly with deterioration of global function and adverse outcomes, highlighting the remote tissue as a mechanistically and therapeutically relevant substrate in post-MI heart failure (6).

Cardiac fibroblasts (CFs) are the central orchestrators of the fibrotic remodelling process. In addition to generating and remodeling extracellular matrix (ECM), activated fibroblasts regulate growth factor bioavailability and mechano-signaling within the myocardial interstitium (4, 7). Prior work from our laboratory demonstrated that CFs isolated from the remote myocardium after MI exhibit proteomic signatures that closely resemble those of infarct fibroblasts, suggesting that a “scar-like” fibroblast program extends into regions not directly exposed to ischemia (4). These findings align with cardiac magnetic resonance studies using native T1 mapping and extracellular volume quantification, which reveal early and diffuse abnormalities in remote myocardium consistent with edema and interstitial fibrosis, even when injury is localized to a discrete infarct territory (3). Complementary histological and molecular analyses in animal models and human hearts further support that remote remodeling involves sustained transcriptional reprogramming, ECM remodeling, and persistent immune signaling (8–10). Together, these observations highlight a major gap in the field: the molecular determinants that drive maladaptive remote ECM remodeling and stiffening after MI remain incompletely defined.

In interrogating fibroblast and myocardial proteomic datasets across infarct and remote regions early after MI, we identified latent transforming growth factor-β binding protein 2 (LTBP2) as one of the most strongly induced ECM-associated proteins in activated CFs and as a prominent secreted candidate in remote myocardium. LTBP2 is a fibrillin-associated microfibrillar glycoprotein known for roles in microfibril assembly and tissue biomechanics, and emerging evidence implicates LTBP2 in fibrotic pathobiology across organs (11–13). In the heart, LTBP2 was identified in an unbiased transcriptomic screen as the gene consistently upregulated in CFs across three mouse strains following isoproterenol challenge, was similarly induced after transverse aortic constriction, and was elevated in plasma and myocardium from patients with heart failure, yet its contribution to the pathogenesis of cardiac fibrosis was left undefined (14). *Ltbp2* knockdown has been reported to attenuate fibrosis in non-ischemic models of cardiac disease, where the effects were attributed to NF-κB signaling and oxidative stress (15, 16), and elevated LTBP2 has been described in human heart failure (17), yet whether LTBP2 contributes to remodeling after ischemic injury, and by what mechanism, has not been established. Notably, although other LTBP family members (LTBP1/3/4) tether latent TGF-β complexes to the ECM to regulate spatially controlled activation, LTBP2 does not covalently bind latent TGF-β. This raises a fundamental mechanistic question: how could an LTBP family member that lacks canonical latent TGF-β binding activity influence the prototypical profibrotic TGF-β signaling axis in the injured heart?

We hypothesized that injury-induced LTBP2 promotes maladaptive remodeling of the remote myocardium by indirectly modulating TGF-β bioavailability. Specifically, we proposed that elevated LTBP2 incorporation into fibrillin-1–rich networks alters the capacity of microfibrils to sequester latent TGF-β complexes carried by other LTBPs, shifting TGF-β compartmentalization toward a more soluble, activatable pool and sustaining profibrotic signaling within viable myocardium (18, 19). To test this, we combined genetic loss-of-function studies with regional functional phenotyping, quantitative histology and biomechanics, remote-tissue proteomics, and gain-of-function experiments using a defined fibrillin-1–binding module of LTBP2.

Here we show that LTBP2 is induced in injury-activated CFs, and elevated in failing human left ventricle across ischemic, non-ischemic, and hypertrophic cardiomyopathy colocalized with FBN1. Global deletion of *Ltbp2* improves post-MI survival, improves regional systolic and diastolic strain in remote segments, and attenuates remote interstitial fibrosis and myocardial stiffening, without altering infarct size, early post-infarct function, or cardiomyocyte hypertrophy. Remote-tissue proteomics identifies TGF-β–responsive fibrotic programs enhanced in wild-type and blunted in *Ltbp2*-deficient hearts. Mechanistically, loss of LTBP2 increases LTBP1 incorporation into fibrillar matrix and retains latent TGF-β there, while a minimal EGF13–16 module of LTBP2 binds fibrillin-1 directly and is sufficient to mobilize latent TGF-β into the activatable pool. These findings identify LTBP2 as a driver of remote myocardial remodeling and define an ECM-centered mechanism, competitive remodeling of microfibril occupancy, linking fibroblast activation to sustained TGF-β signaling and progression toward heart failure after MI.

## Supplemental Methods

### Myocardial Infarction & Mouse Model

All experiments were approved by the University of Toronto’s Institutional Animal Care and Use Committee and conducted in compliance with Canadian Council on Animal Care guidelines and national animal protection laws (Animal Use Protocol: AUP:20011793, and AUP:20011218). *Ltbp2 -/-*mice were provided from Dr. Robert Mechams Lab. Adult C57Bl/6 mice (10–16 weeks old, littermate) were housed under standard 14:10 h light–dark cycles with free access to water and chow. Myocardial infarction (MI) was surgically induced via permanent ligation of the Left Anterior Descending (LAD) coronary artery as described previously (4, 20). Briefly, under anesthesia, a 7-0 Prolene suture was passed beneath the LAD approximately 2 mm below the left atrium and tied to occlude blood flow. Sham-operated mice underwent identical procedures, including suture placement, but without ligation. Following surgery, all animals recovered for at least 7 days before subsequent analysis. Both male and female mice were used for all analysis aside from echocardiographic assessment.

### Echocardiography

Cardiac structure and function were assessed in sham-operated and post-MI mice at 7 and 28 days using a high-frequency ultrasound system (Vevo 3100, FUJIFILM VisualSonics) with a 25–30 MHz linear-array transducer (MX400), as previously described (21). Mice were lightly anesthetized with isoflurane (induction 3–4%, maintenance 1–1.5% in oxygen), and body temperature, heart rate, and respiration were continuously monitored via a heated ECG platform. Hair was removed from the chest with depilatory cream, and pre-warmed ultrasound gel was applied. All acquisition and offline analyses were performed by an operator blinded to genotype and surgical group.

Parasternal long- and short-axis B-mode and M-mode images were acquired to quantify left ventricular (LV) wall thicknesses, end-diastolic and end-systolic internal diameters (LVEDD, LVESD), fractional shortening (FS), fractional area change (FAC); heart rate was recorded from the simultaneous ECG. To resolve regional systolic function, ECG-gated kilohertz visualization (EKV) reconstructions of the LV long axis were acquired at an effective temporal resolution of 1,000 frames/s, and the endocardial border was traced to calculate FAC. For segmental analysis, the LV long axis was divided into basal, mid, and apical thirds, and M-mode traces were recorded at each level to measure wall thickness and chamber dimensions at end-diastole and peak systole; segment-specific FS was derived from the corresponding LVEDD and LVESD. Diastolic function was assessed by pulsed-wave Doppler of trans-mitral inflow to measure early (E) and late (A) filling velocities, and by pulsed-wave tissue Doppler of the septal mitral annulus to measure early diastolic annular velocity (e′); the E/A and E/e′ ratios were calculated. Pulmonary venous flow was additionally recorded by pulsed-wave Doppler. LV myocardial deformation was quantified by two-dimensional speckle-tracking echocardiography (VevoStrain, FUJIFILM VisualSonics) from B-mode long-axis cines. Two to three consecutive cardiac cycles of comparable quality were selected per animal; the LV endocardial border was traced manually in a single frame and propagated automatically across the cardiac cycle by the software, with tracking quality visually verified and manually corrected where needed. Peak longitudinal strain and strain rate were averaged across the analyzed cycles both globally and for individual (basal, mid, apical) segments. In addition, instantaneous strain and strain-rate were exported frame-by-frame across a representative cardiac cycle to generate continuous deformation-versus-time curves for group-level comparison.

### Histological Preparation and Picrosirius Red Staining

Mouse hearts were carefully excised and rinsed with chilled DPBS (Sigma, D8662-500ML) before immersion in 4% PFA for a two-hour fixation period. Tissues were subsequently dehydrated overnight in a 30% sucrose solution at 4°C until complete tissue submersion was achieved. Following dehydration, the hearts were placed in a 50:50 solution of 30% sucrose and optimal cutting temperature (OCT) compound (Fisher Healthcare, 23-730-571) and incubated for 30 minutes while rotating at 4°C.To prepare for cryosection, the hearts were embedded in OCT using cryomolds and rapidly frozen through immersion in 2-methylbutane with dry ice, then stored at -80°C until sectioning. Cryosections with a thickness of 10 micrometres were collected and affixed to charged slides (VWR, 5027-CA48311-703). Assessment of cardiac fibrosis was conducted through Picrosirius Red staining, following established protocols (22). Sections were further fixed in 4% PFA for 10 minutes at room temperature and washed with DPBS for 5 minutes. A FastGreen/Picrosirius Red dye solution (Chondrex, 9046) was applied to each section, fully immersing them, and incubated at room temperature for 45 minutes. Subsequently, sections were washed with distilled H2O and dehydrated sequentially in concentrated ethanol (50%, 70%, 95%, 100%) for 45 seconds each, followed by 90 seconds in xylene at room temperature. The completed slides were mounted using Cytoseal60 (Thermo Scientific, 23244256) and imaged using a slide scanner (Aperio AT2). Quantification of percent fibrosis was performed using NIH ImageJ software. Researchers acquiring images and quantifying percent fibrosis were blinded to condition and genotype of the sections.

### RNAscope Staining and Analysis

Multiplexed in situ hybridization was performed on cardiac cryosections using the RNAscope Multiplex Fluorescent v2 Assay (Advanced Cell Diagnostics, Newark, CA) according to the manufacturer’s instructions, with minor optimizations for cardiac tissue as described above. Sections were hybridized with a panel of 11 target probes, of which Ltbp2 and Pdgfra are reported here. Because the assay accommodates three fluorescent channels per round, probes were applied in four sequential rounds of three. Following each round, nuclei were counterstained with DAPI and whole-slide images were acquired on a Zeiss AxioScan fluorescence slide scanner (Zeiss, Oberkochen, Germany) using identical exposure and gain settings across all slides. Probes and fluorophores were then stripped according to the manufacturer’s protocol, and the next set of three probes was applied to the same section. This cycle was repeated until all 11 probes had been imaged.

Image analysis was performed in HALO (Indica Labs, Albuquerque, NM) at the Advanced Optical Microscopy Facility, Princess Margaret Cancer Research Tower. The four imaging rounds were registered to one another using the DAPI channel, allowing all 11 probe signals to be visualized within the same cells. Signal thresholds were set per channel to distinguish specific puncta from background sections as a reference and were then applied uniformly across all slides. Cells were classified by the combination of probes detected. For this study, Ltbp2⁺/Pdgfra⁺ cells were identified as fibroblasts expressing Ltbp2 and rendered as spatial overlays in HALO for display in Figure 1D.

**Figure 1.**
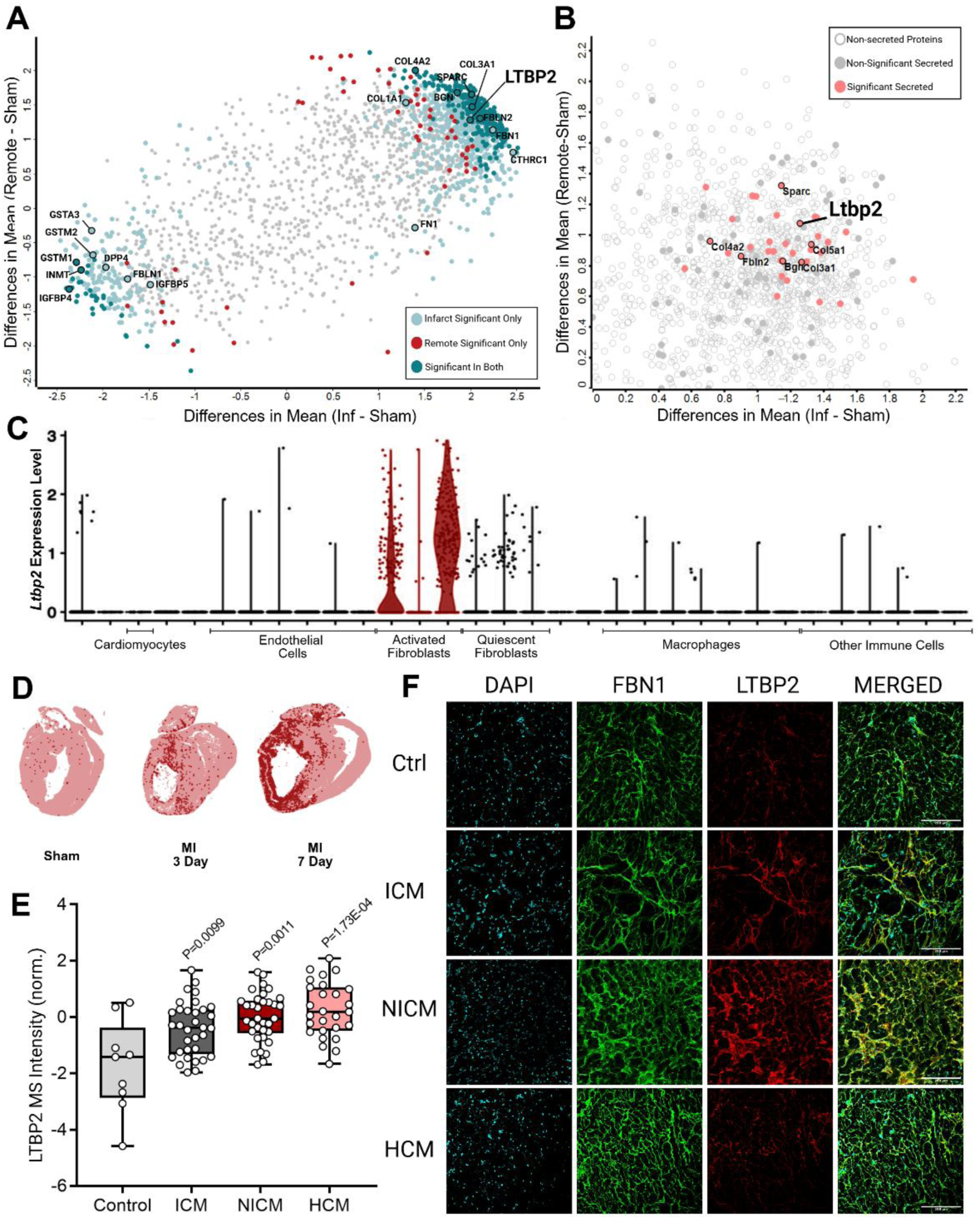
LTBP2 is upregulated in injury-activated cardiac fibroblasts of infarct and remote myocardium after myocardial infarction and is increased in failing human left ventricle. (A) Scatter plot of ventricular cardiac fibroblast proteins commonly identified by mass spectrometry at 7 days after left anterior descending coronary artery ligation, plotting the difference in mean z-scored intensity for infarct versus sham (x-axis) against remote versus sham (y-axis). Points are coloured by the comparison in which significance was reached: infarct only (light blue), remote only (red), or both regions (teal); grey denotes proteins not significant in either comparison (adjusted P < 0.01). Selected extracellular matrix and fibroblast-associated proteins are labelled, with LTBP2 among those significantly increased in both regions. (B) Proteins with positive mean z-scores in A replotted to distinguish secreted from intracellular candidates. Open circles, non-secreted proteins; grey filled circles, secreted proteins not reaching significance; pink filled circles, secreted proteins significantly increased in remote versus sham fibroblasts (adjusted P < 0.05). *Ltbp2* is among the most prominently upregulated secreted matrix proteins. Raw proteomic data from Shah et al. (C) De novo analysis of published single-cell RNA sequencing data from the infarcted mouse heart (Farbehi et al. 2019) showing *Ltbp2* expression across annotated cardiac cell populations. Expression is largely restricted to injury-induced, activated fibroblast clusters (red) and is minimal in quiescent fibroblasts, cardiomyocytes, endothelial cells, macrophages, and other immune populations. (D) RNAscope in situ hybridization for *Ltbp2* on whole-heart sections from sham-operated mice and mice at 7 days after myocardial infarction. Each red punctum represents a single *Ltbp2^+^*/*Pdgfra*^+^ cells marking for *Ltbp2* expressing fibroblasts. (E) Normalized LTBP2 mass spectrometry intensity in left ventricular tissue from non-failing controls and patients with ischemic (ICM), non-ischemic (NICM), or hypertrophic (HCM) cardiomyopathy. Each point represents one patient; n = 9 control, 33 ICM, 34 NICM, 27 HCM. P values are shown for each cardiomyopathy group versus control and were corrected for multiple comparisons using the Benjamini-Hochberg false discovery rate procedure. (F) Representative immunofluorescence staining of human left ventricular myocardium for FBN1 (green), LTBP2 (red), and nuclei (DAPI, cyan), with the merged channel at right. LTBP2 deposition is increased in ICM and NICM myocardium relative to control and overlaps FBN1-positive microfibrillar structures. Images are representative of n = 1 for all groups are presented for qualitative comparison; staining was not quantified. Scale bars, 200 μm.

### Tissue Preparation for Mass Spectrometry

Remote heart tissue was homogenized in freshly prepared 8M urea using a combination of Dounce homogenization and sonication (40% intensity for 30 seconds) to ensure mechanical shearing. Following homogenization, cellular debris was removed via sequential centrifugation at 20,000 RCF at 4°C in a microcentrifuge until no visible pellet remained. Total protein content in the clarified lysate was quantified using a Coomassie/Bradford assay, with urea diluted to <1M to ensure assay compatibility. For trypsin digestion, 10–50 µg of protein was allotted, adjusted to equal volumes with 50 mM ammonium bicarbonate (ABC), and reduced with 2.5 mM DTT (37°C, 60 min). Alkylation was performed in the dark using 5 mM iodoacetamide for 30 min. Samples were diluted 10-fold in 50 mM ABC, digested overnight at 37°C with sequencing-grade trypsin (1:20 enzyme-to-protein ratio), and acidified with formic acid (1% final concentration) to terminate the reaction. Residual lysate was stored at -80°C as backup.

### Liquid Chromatography Mass Spectrometry Analysis

Peptides were desalted, loaded onto C18 Evotips (Evosep Biosystems, EV-2001) at ∼1 μg per sample, and analyzed on an Evosep One LC system (EV-1000) coupled to a Q Exactive Plus mass spectrometer (Thermo Fisher Scientific) as described previously (23). Data-independent acquisition was performed in positive ion mode across 400–971 *m/z*, with each cycle comprising one MS1 scan (70,000 resolution; AGC target 3 × 10⁶; maximum injection time 200 ms) followed by 30 DIA MS/MS scans (17,500 resolution; AGC target 5 × 10⁵; normalized collision energy 27) using 20 *m/z* isolation windows with 1 *m/z* overlap. Raw files were processed in DIA-NN v1.8.1 using a spectral library–free search against the *in silico*–predicted mouse proteome (UniProt UP000000589, February 2024 release; canonical and isoform sequences, ∼55,000 entries). Trypsin specificity allowed one missed cleavage, with carbamidomethylation of cysteine as a fixed modification and methionine oxidation as variable; precursor FDR was controlled at 1%. Scan window, retention time alignment, and mass accuracy (12–30 ppm) were inferred automatically. Match-between-runs, unrelated runs, and isotopologue options were enabled, with RT- and signal-dependent cross-run normalization and robust LC (high precision) quantification.

Peptide and protein intensities were log2-transformed prior to downstream analysis. Proteins detected in at least 70% of samples in one experimental group were retained for analysis, and missing values were imputed from a normal distribution (width = 0.30, downshift = 1.80). Differential expression analysis was performed using unpaired two-sample Student’s t-tests in Perseus (v2.0.10.0), with significance defined at a false discovery rate (FDR) < 0.05. Principal component analysis (PCA) and hierarchical clustering (Euclidean distance, k-means preprocessing with 300 clusters and 10 iterations) were used for global visualization of proteomic variation between groups.

### Atomic Force Microscopy

Left ventricular tissue from wild-type (WT: n = 4 sham, n = 4 infarct/remote) and LTBP2 knockout (KO: n= 4 sham, n = 4–5 infarct/remote) mice was analyzed 4 weeks after LAD ligation or sham surgery. Hearts were excised, rinsed in ice-cold Ca²⁺/Mg²⁺-supplemented Dulbecco’s phosphate-buffered saline (DPBS), and sectioned into 1 mm-thick longitudinal slices using a vibratome to maximize infarct and remote tissue inclusion as outlined previously (24). Freshly sectioned samples were immobilized on cover glass with adhesive glue and maintained in DPBS at 4°C prior to atomic force microscopy (AFM).

Mechanical testing was performed using an MLCT-BIO probe A (nominal spring constant: 0.07 N/m), calibrated thermally before each session. All tips were calibrated on glass with contact-based and contact free methods, probes with sensitivity >20 nm/V and indentation rates <2 µm/s were employed to minimize viscoelastic effects and ensure quasistatic elasticity measurements (25,26). To mitigate batch variability, WT and KO samples were interleaved during testing, with operators blinded to genotype and tissue region (sham, infarct, remote). For each sample, 15–35 force-indentation curves were acquired across an 8 × 8 µm grid with constant extend speed, set point, and z-length across samples. Elastic modulus was derived by fitting retraction curves to a Hertz quadratic pyramidal contact model. Technical replicates per sample and biological replicates per group (n ≥ 4) were aggregated for statistical comparisons.

### Immunofluorescence Microscopy

Tissue samples embedded in optimal cutting temperature compound were sectioned and fixed in 4% paraformaldehyde (PFA; Sigma, Cat# P6148) in calcium/magnesium-free PBS for 10 min at room temperature. Sections were permeabilized with 0.1% Triton X-100 (BioShop, Cat# TRX777) in PBS (10 min), then blocked for 1 h at room temperature with 5% bovine serum albumin (BSA; BioShop, Cat# ALB001) in PBS. Primary antibodies, LTBP2 (From Dr. Nakamura) and FBN1 (From Dr. Nakamura) (18), were diluted in PBS/5% goat serum and incubated overnight at 4°C. After PBS washes, sections requiring secondary detection were incubated for 1 hour at room temperature in the dark with goat anti-rabbit IgG Alexa Fluor 488 (Thermo Scientific, Cat# A11034, 1:1000) and goat-anti chicken IgG Alexa Fluor 564 (Invitrogen Cat # A-21449). Nuclei were counterstained with 2 μg/mL DAPI (10 min, room temperature). Imaging was performed using an Olympus FV3000 laser-scanning confocal microscope, with consistent laser power, gain, and pinhole settings across samples. Cell culture samples were prepared similarly to tissue sections on Mattek Plates.

### Wheat Germ Agglutinin Staining and Analysis Protocol

Following fixation and blocking, cardiac cryosections were incubated with CF488A-conjugated wheat germ agglutinin (WGA) (Biotium, Cat# 29022-1) at 5 μg/mL for 30 minutes at room temperature to delineate cardiomyocyte membranes. Nuclei were counterstained with 2 μg/mL DAPI for 10 minutes at room temperature, and sections were mounted as described above.

Images were acquired from the remote region of the left ventricle at 20× magnification on an Olympus FV3000 laser-scanning confocal microscope, with identical laser power, gain, and pinhole settings across all samples. Cardiomyocyte cross-sectional area was measured manually in ImageJ/Fiji by tracing the WGA-delineated membrane boundary. Only transversely sectioned cardiomyocytes with a circular profile and a centrally located nucleus were included, to exclude obliquely sectioned cells. Approximately 100 cardiomyocytes were measured averaged across 2 fields analyzed per animal, and the per-animal mean cross-sectional area used for statistical comparison. Image acquisition and analysis were performed by an investigator blinded to genotype and surgical group. At least 4 biological replicates were used per group.

### Primary Cardiac Fibroblast Isolation

Our CF isolation protocol was adapted from a strategy used by Pinto and colleagues (4, 27). Specifically, the whole heart of an adult C57Bl/6 mouse was excised, and the atria and atrioventricular valves were surgically removed. The ventricles and septum were minced into 1 mm pieces and digested with 2 mg/mL collagenase type II (Worthington Cat#: LS004176) and 2 mg/mL dispase II (Sigma Cat#: D4693) in Dulbecco’s phosphate-buffered saline (dPBS) supplemented with 0.9 mmol/L CaCl2 for 45 min at 37 °C with periodic mixing every 15 min. The digest was then filtered through a 70 μm mesh into 25mL of 1× dPBS. Next, the digest was centrifuged for 7 min at 500g at 4 °C. Resulting pellets were resuspended in fibroblast media, containing 1% l-glutamine (Thermo Cat#: 25030-081), 1% penicillin–streptomycin (Thermo Cat#: 15140-122), and 10% fetal bovine serum (Thermo Cat# 10437-028) in Dulbecco’s modified Eagle’s media (Thermo Cat#: 11965-092), and plated for 2 h on a standard tissue culture-treated Petri dish. Following the 2 h prelate, old media was aspirated and replaced with fresh fibroblast media. The media was subsequently replaced every other day until the end of the experiment.

### Gene Expression Quantification by Reverse Transcript – Quantitative Polymerase Chain Reaction

Expression of *Ltbp1–4* and *Fbn1* was assessed in primary cardiac fibroblast cultures by reverse transcription quantitative polymerase chain reaction (RT-qPCR). Total RNA was isolated using the RNeasy Mini Kit (Qiagen, 74104) according to the manufacturer’s instructions, including on-column DNase I digestion to remove genomic DNA. RNA concentration and purity were assessed by spectrophotometry (NanoDrop; A260/280 and A260/230 ratios), 500 ng of RNA per sample was reverse transcribed using SuperScript III Reverse Transcriptase (Life Technologies, 18080044) with random hexamers following the manufacturer’s protocol. Reverse transcriptase–negative controls were prepared in parallel to confirm the absence of genomic DNA amplification.

cDNA was diluted to 2 ng/μL, and reactions were assembled with Power SYBR Green Master Mix (Applied Biosystems, 4367659). Amplification was performed on a BioRad CFX384 using the cycling conditions provided by SYBR Green protocol. A melt curve was acquired after each run to confirm single-product amplification. All reactions were run in technical triplicate with at least four biological replicates, and no-template controls were included on every plate. Primer sequences are provided in major resource tables. Relative gene expression was calculated by the 2^−ΔΔCt method with *Gapdh* as the reference gene, and results are expressed as fold change relative to [day 1 cultures / BSA vehicle control] as indicated in the figure legends.

### Silicone Substrate for Cell Plating

Silicone substrates of defined stiffness were prepared using the Sylgard 527 A&B silicone dielectric gel kit (Dow Corning, 1696742). Components A and B were combined in 50 mL conical tubes at ratios calculated to yield target elastic moduli of 15, 50, and 70 kPa according to the relationship described previously (28). Mixtures were degassed under vacuum for 15 minutes, then dispensed slowly into plastic culture dishes at 105 μL/cm² (0.5 mL per well for 12-well plates, 1 mL per well for 6-well plates, 6 mL per 10 cm dish, 15 mL per 15 cm dish). Plates were covered with lids set at an angle and allowed to degas for a further 30 minutes at room temperature before curing at 75°C for 48 hours on a paper towel underlay to prevent deformation of the plastic.

Cured gels were functionalized with collagen to permit cell attachment. Gel surfaces were plasma-treated at high power for 20 seconds and immediately covered with 1% (v/v) 3-aminopropyltriethoxysilane (APTES; Sigma-Aldrich, 440140) in ddH₂O, scaled to vessel size (2 mL per well for 12-well plates, 4 mL per well for 6-well plates, 25 mL per 10 cm dish, 75 mL per 15 cm dish), and incubated at 75°C for 90 minutes. APTES was aspirated and gels were washed in ddH₂O for 15 minutes, incubated in 0.1% paraformaldehyde in Ca²⁺/Mg²⁺-free PBS for 20 minutes at room temperature with gentle agitation, and washed again in ddH₂O for 15 minutes. Plates were then sterilized under ultraviolet light in a biosafety cabinet for 20–30 minutes with lids removed. Gels were coated with 100 μg/mL PureCol bovine collagen I (Sigma-Aldrich, Cat# 804622) in Ca²⁺/Mg²⁺-free PBS, scaled to vessel size (1.5 mL per well for 12-well plates, 3 mL per well for 6-well plates, 15 mL per 10 cm dish, 50 mL per 15 cm dish), and incubated overnight at 4°C. Excess collagen solution was washed off with Ca²⁺/Mg²⁺-free PBS immediately before cell seeding. Standard tissue culture–treated polystyrene (>1,000 kPa), coated identically with collagen, served as the stiff control condition.

### TGF-β Reporter Assays

To quantify bioactive TGF-β in cardiac fibroblast (CF) cultures, we used the transformed mink lung epithelial cell reporter line (TMLC) harboring a PAI-1 promoter–driven luciferase construct (29). CFs were plated at equal density in 24-well plates and maintained for 5, 7, 9, 11, 21, or 28 days post-plating, with media changes every 48 hours. Twenty-four hours prior to collection, cells were switched to DMEM containing 1% FBS that had been heat-inactivated (80°C, 2 hours) to minimize exogenous TGF-β contribution while permitting accumulation of fibroblast-derived TGF-β. Conditioned media (CM) was collected and divided for assessment of: active TGF-β, measured directly from CM, and total TGF-β, determined following heat activation (80°C for 10 minutes) to convert latent TGF-β to its active form. Following CM collection, cultures were washed with DPBS containing Ca²⁺/Mg²⁺ and decellularized to isolate cell-derived extracellular matrix (ECM), as previously described (30). To extract matrix-bound latent TGF-β, ECM was subjected to acid activation using glycine-HCl (pH 2.0), followed by neutralization with Tris-HCl (pH 9.5), consistent with established protocols for liberating latent TGF-β from the matrix. TMLC reporter cells were plated in 96-well plates and serum-reduced (1% heat-inactivated FBS) 8 hours prior to treatment. CM (active and total fractions) or acid-activated ECM extracts were applied to TMLC cells for 24 hours. Luciferase activity was measured using the Luciferase Assay System (Promega, #E1500) on a Berthold LB Centro plate reader. For luminescence detection, luciferase substrate (1:3 dilution) was injected into each well, shaken for 3 seconds, incubated for 10 seconds, and luminescence recorded for 10 seconds per well. Standard curves were generated for each assay plate using recombinant human TGF-β1 across 0–3,330 pg/mL and fitted by linear regression; curves with r² < 0.98 were not used for interpolation. Conditioned media were diluted 1:2 and acid-activated ECM extracts 1:6 before application to TMLC cells, such that all interpolated values fell within the range of the standard curve. Reported concentrations are corrected for dilution.

To investigate the effects of exogenous LTBP2 on TGF-β signaling, primary fibroblasts were treated with purified His-FLAG-LTBP2 at a final sample concentration of 1.5 µM, for either 20 mins or 24 hours. Heat-denatured His-FLAG-LTBP2, prepared by boiling the purified protein product for 10 mins, was included as a control. Following treatment, TGF-β signaling was assessed using the reporter assay described above.

### Integrin αV Inhibition in Cardiac Fibroblasts Cell Culture

Integrin inhibition experiments were performed as part of the primary CFs culture time course. CWHM-12, a small-molecule pan-αv integrin antagonist (gift from Dr. Boris Hinz, University of Toronto), and its inactive enantiomer CWHM-96, used as vehicle control, were prepared as 10 mM stock solutions in cell culture dimethyl sulfoxide (DMSO) and stored at -20°C. Stocks were diluted in culture medium to a final working concentration of 5 μM.

Wild-type and *Ltbp2* knockout CFs were plated at equal density and maintained as described above. At days 7, 9, and 11 after plating, cultures were treated with 5 μM CWHM-12 or 5 μM CWHM-96 for 24 hours prior to collection. Compound was added concurrently with the switch to DMEM containing 1% heat-inactivated FBS (80°C, 2 hours) used for conditioned media collection, so that inhibitor exposure spanned the entire accumulation window. Conditioned media and cell-derived extracellular matrix were subsequently collected, and active, total, and matrix-associated TGF-β were quantified by TMLC reporter assay exactly as described above.

### LTBP2 and FBN1 Construct Design

Recombinant expression constructs were generated by GenScript and verified by whole-plasmid sequencing. For lentiviral expression, a construct encoding human LTBP2 epidermal growth factor-like domains 13–16 (EGF13–16; amino acids 1217–1410) was generated in the pLVV-EF1α-PGK-Puro-P2A-EGFP lentiviral expression backbone. The construct contains an N-terminal preprotrypsin signal peptide (MSALLILALVGAAVA), followed by a FLAG epitope tag (DYKDDDDK), a flexible glycine-serine linker (GGGGS), and the LTBP2 coding sequence. The corresponding empty lentiviral backbone, lacking the LTBP2 insert, was used as a control.

For protein purification, constructs encoding human LTBP2 EGF13–16 (amino acids 1217–1410) and human FBN1 (amino acids 19–489) were generated in the pcDNA3.1 mammalian expression vector. Both constructs contained an N-terminal preprotrypsin signal peptide followed by a hexahistidine (6xHis) affinity tag, a tobacco etch virus (TEV) protease cleavage site (ENLYFQG), and a flexible glycine-serine linker (GGGGS). The LTBP2 construct contains a FLAG epitope tag (DYKDDDDK), whereas the FBN1 construct contains a MYC epitope tag (EQKLISEEDL), each separated from the corresponding protein sequence by an additional GGGGS linker.

### Cell Culture and Recombinant Protein Purification

Human embryonic kidney cells (HEK-293T; ATCC, CRL-3216,) were grown and maintained in 150 cm2 cell culture dishes using Dulbecco’s Modified Eagle Media (DMEM; Wisent, #319-005-CL) supplemented with 10% fetal bovine serum (FBS), 0.1 mM MEM non-essential amino acids (Gibco, #11140-050) and 1% penicillin-streptomycin.

Cells were transfected with 22 µg of the transgene plasmid (His-FLAG-LTBP2 or His-MYC-FBN1) using the polyethyleneimine transfection reagent (PEI; Polysciences Inc., #23966). At approximately 16-18 hours post-transfection, culture medium was replaced with serum-free DMEM supplemented with 1% penicillin-streptomycin. Media containing secreted recombinant proteins was collected 48-72 hours post-transfection and supplemented with EDTA-free protease inhibitor cocktail (Sigma, #11836170001). Collected media was clarified by sequential centrifugation at 500 x g for 5 mins followed by 4000 x g for 10 mins at 4°C. Clarified supernatants were adjusted to a final concentration of 10 mM imidazole and 300 mM NaCl prior to purification. The media was then incubated with Ni-NTA agarose resin (Qiagen, #30210) via batch binding for 2 hours rotating at 4°C. The resin was pre-equilibrated in binding buffer containing 50 mM Tris, 300 mM NaCl, and 10 mM imidazole (buffer A; pH 7.8-8.0). Following protein binding, the resin was washed with binding buffer A, followed by washes in a buffer containing 50 mM Tris, 300 mM NaCl, and 20 mM imidazole (buffer B; pH 7.8-8.0). Bound proteins were eluted using 50 mM Tris, 300 mM NaCl, and 250 mM imidazole (buffer C; pH 7.8-8.0). Eluted proteins were subsequently buffer exchanged into 20 mM HEPES, 50 mM NaCl and 2 mM CaCl2 (pH 7.5) using centrifugal desalting columns, with a norminal molecular weight cut-off of 7 kDa (ThermoFisher Scientific, #89891). Total protein concentration was determined using a BCA Protein Assay Kit (ThermoFisher Scientific, #23235).

### Lentiviral Transduction of Isolated Murine Fibroblasts

For lentiviral production, HEK-293T cells were transfected in 150 cm^2^ culture dishes with 12 µg transgene plasmind (pLEX-GFP or pLEX-hLTBP2), 6 µg packaging plasmid (pSPAX2) and 3 µg envelope plasmid (pMD2.G) using polyethyleneimine. 72 hours post-transfection, lentivirus-containing culture media was collected and centrifuged at 1000 x *g* for 5 min at 4 °C, to remove cellular debris. Supernatants were subsequently filtered through a 0.45 µm membrane and concentrated using centrifugal filter units with a nominal molecular weight cutoff of 50 kDa (Millipore Sigma, UFC9050). Five days after seeding, primary fibroblasts were transduced with concentrated lentiviral supernatants supplemented with 10 µg/mL polybrene (Millipore Sigma, #TR-1002) for 21 hours at 37°C. Lentiviral transduction was performed using a plate-to-plate ratio, whereby viral supernatant generated from a single 150 cm^2^ culture dish was used to transduce one well of a 24-well plate. Cells were harvested 72 hours post-transduction

### Co-immunoprecipitation

Anti-FLAG M2 Affinity Gel (Sigma-Aldrich) was blocked with 0.1% bovine serum albumin (BSA) in immunoprecipitation (IP) buffer containing 20 mM HEPES, 150 mM NaCl, 2 mM CaCl₂, 0.05% Triton X-100, and 5% glycerol (pH 7.4) supplemented with EDTA-free protease inhibitor for 30 mins at 4°C. For FLAG affinity pull-down, blocked beads were incubated with 4.7 µg of total protein from the purified His-FLAG-LTBP2 preparation (estimated final concentration of 1 µM) for 2 hours at 4°C with rotation to facilitate bait protein immobilization. Following incubation, beads were washed twice with IP buffer before bead bound His-FLAG-LTBP2 was subsequently incubated with 11 µg of total protein from the purified His-MYC-FBN1 preparation (estimated final concentration of 1 µM), overnight at 4°C with rotation. Following incubation, beads were washed three times with IP buffer to remove unbound proteins. Bound protein complexes were eluted and denatured in Laemmli sample buffer (2X) supplemented with 10 mM DTT, boiled for 5 mins and resolved by SDS-PAGE.

### Immunoblotting

Laemmli-solubilized samples were run on an SDS-PAGE gel, followed by transfer onto a nitrocellulose membrane and blocking with 5% (w/v) skim milk. Following blocking, the membrane was probed with appropriate primary antibodies (Cell Signalling Technologies, rabbit polyclonal FLAG 1:1000 and rabbit polyclonal MYC 1:1000) overnight at 4 °C. Membranes were then incubated with a horseradish peroxidase conjugated secondary antibody (Promega, goat anti-rabbit HRP 1:5000) the next day for 1 hour at room temperature. Chemi-luminescence signal was visualized using a Pierce ECL Western Blotting Substrate (Thermo Fisher Scientific, #32106).

### Human Myocardial Tissue Acquisition and Processing

This work was conducted with the approval of the University of Toronto Human Participants Ethics Board (REB# 45647) and University Health Network Research Ethics Board (REB# 15-8781, 18-6012). All patients with end-stage HF prior to heart transplantation provided informed written consent. Control tissues were obtained through the Trillium Gift of Life Network following standard protocols for donation after brain death, from organs deemed unacceptable for donation, with normal echocardiograms and coronary angiograms. Biospecimens from the left ventricular free wall were collected, immediately snap frozen in liquid nitrogen, and stored at -80°C until use. Samples prepared for immunofluorescences were embedding in OCT compound and sectioned at 5 um thickness using a cryostat and stained and imaged similar to mouse cardiac tissue as described above.

### Failing Heart Proteome Analysis

Publicly available liquid chromatography tandem mass spectrometry (LC-MS/MS) datasets were analysed from published studies assessing the global cardiac proteome from end-stage human heart failure. These datasets included Hamed et al. (PXD068439) (31). Protein expression levels are shown as log-transformed, width adjustment normalized and imputed from a normal distribution.

### Cell Culture ECM Isolation for Mass Spectrometry

Wild-type and *Ltbp2* knockout primary CFs were plated at equal density in 24-well plate and maintained for 7, 11, or 21 days with media changed every 48 hours. Cultures were washed three times with Ca²⁺/Mg²⁺-supplemented DPBS to remove residual media, then decellularized in 0.1% (v/v) Triton X-100 (BioShop, TRX777) with 20 mM ammonium hydroxide in DPBS for 10 minutes at 37°C, following established protocols for preserving cell-derived matrix (4). Decellularized matrix was washed three times in Ca²⁺/Mg²⁺-supplemented DPBS to remove cellular debris and residual detergent. Preservation of matrix architecture, and removal of nucleic material after decellularization was confirmed in parallel wells by immunofluorescence for LTBP2 and FBN1 and DAPI [Supplementary Figure 6].

Matrix was solubilized in freshly prepared 8 M urea and recovered by mechanical scraping from the culture surface. Samples were sonicated at 40% amplitude for 30 seconds to complete solubilization and shear residual nucleic acid, then clarified by sequential centrifugation at 20,000 × g at 4°C until no visible pellet remained. Protein concentration was determined by Coomassie/Bradford assay with urea diluted to <1 M for assay compatibility. Reduction, alkylation, tryptic digestion, desalting, and liquid chromatography–mass spectrometry acquisition were performed exactly as described above for myocardial tissue. n = 5 independent wild-type and 4 knockout cultures per timepoint.

### Skeleton Length Analysis for LTBP1 staining in Wt and KO Primary Cardiac Fibroblasts

Fibrillar deposition of LTBP1 and LTBP2 in wild-type and *Ltbp2* knockout primary cardiac fibroblast cultures was quantified from immunofluorescence images acquired at days 7,9,11, and 21 after plating, following the unbiased extracellular matrix fiber quantification protocol developed by the Reinhardt group for fibrillin and related microfibrillar fiber systems (32). All images were acquired on an Olympus FV3000 confocal microscope at the same magnification using identical laser power, gain, and pinhole settings, and exported as lossless 8-bit TIFF files. All processing steps were applied identically across genotypes, timepoints, and channels.

Analysis was performed in ImageJ/Fiji. Single-channel images were converted to 8-bit grayscale. Images were thresholded using Image > Adjust > Threshold with “Default,” “B&W,” and “Dark background” selected; the threshold maximum was set to 255 and the minimum set to the average optimized value determined across a subset of images, then held constant for every image analyzed so that measurements were directly comparable between conditions. Background subtraction was not applied. Noise from small non-fibrillar particles was removed by applying Process > Noise > Despeckle three times. Thresholded images were skeletonized using Plugins > Skeleton > Skeletonize (2D/3D), reducing fibrillar structures to single-pixel-wide representations, and analyzed with Analyze > Skeleton > Analyze Skeleton (2D/3D) with the prune cycle method set to “none” and detailed information enabled. Total skeleton length, number of branches, average branch length, and number of junctions were extracted from the Results and Branch Information outputs. Analysis was performed on n = 4–5 independent fibroblast isolations per genotype and timepoint. Per-image values were averaged to give a single value per isolation, and statistical comparisons were performed on these biological replicates by an investigator blinded to genotype.

### Statistical Analysis

Statistical analyses were performed using GraphPad Prism (GraphPad Software, San Diego, CA) and Perseus. Data are presented as mean ± SEM unless otherwise indicated. For comparisons involving a single independent variable, one-way ANOVA was used; for experiments with two independent variables two-way ANOVA was applied. When significant main or interaction effects were detected, Tukey post hoc testing was performed to correct for multiple comparisons. For proteomic analyses, differential protein abundance was assessed using two-tailed Student’s t-tests, and results were visualized using volcano plots. Multiple testing correction was performed using a false discovery rate (FDR) threshold of 0.05 to define statistical significance. The specific statistical tests used, exact n values (biological replicates), and significance levels are provided in the corresponding figure legends. A two-sided P < 0.05 was considered statistically significant.

## Results

### Injury-induced cardiac fibroblasts upregulate and deposit LTBP2 in infarct and remote viable myocardium after myocardial infarction

To identify ECM regulators associated with post-MI remodeling, we interrogated a proteomic dataset previously generated in our laboratory from CFs isolated from infarct and remote myocardial regions at 7 days post-MI (4). Among the proteins most strongly upregulated in both regions, LTBP2 emerged as a significantly increased ECM-associated candidate (Figure 1A). Restricting the analysis to proteins with predicted secretion signals confirmed enrichment of fibrotic, matrisomal, and ECM-associated proteins in post-MI CFs, with LTBP2 among the most prominently upregulated secreted ECM proteins in remote fibroblasts (Figure 1B). To define the cellular source of *Ltbp2*, we mined published single-cell RNA sequencing data from the infarcted mouse heart (7). *Ltbp2* expression was largely confined to injury-induced, activated fibroblast populations and was minimal in quiescent fibroblasts, cardiomyocytes, endothelial cells, and immune cells (Figure 1C). Asking whether this induction reflected fibroblast activation more broadly, we found that *Ltbp2* was significantly upregulated in primary CFs by both TGF-β stimulation and increased substrate stiffness (Supplementary Figure 1A–B), with corresponding accumulation of LTBP2 and FBN1 in fibrillar ECM structures on stiffer substrates (Supplementary Figure 1C). *Ltbp2* induction is therefore a shared feature of activated CF states triggered by distinct profibrotic cues.

Consistent with these findings, multiplexed RNAscope in situ hybridization identified *Ltbp2*-expressing CFs (*Pdgfra*⁺) within the infarct and border zone by 3 days post-MI, with these cells extending into remote, non-infarcted myocardium by day 7 (Figure 1D). At the protein level, LTBP2 accumulated markedly within the infarcted apex and was also detectable in remote basal myocardium at 7 days post-MI, where it overlapped FBN1-positive structures (Supplementary Figure 1D), consistent with incorporation into fibrillin-rich microfibrillar ECM. To determine whether these observations extend to human disease, we examined LTBP2 in left ventricular tissue from patients with ischemic (ICM, n = 33), non-ischemic (NICM, n = 34), or hypertrophic (HCM, n = 27) cardiomyopathy alongside non-failing controls (n = 9). LTBP2 abundance measured by mass spectrometry was significantly increased in all three disease groups relative to controls (Figure 1E), and immunofluorescence staining of representative sections showed LTBP2 deposition overlapping FBN1-rich microfibrillar structures in diseased myocardium (Figure 1F). Together, these findings identify LTBP2 as an injury-induced, fibroblast-derived ECM protein that is deposited within fibrillin-rich matrix in both infarct and remote myocardium early after MI, is responsive to multiple fibroblast-activating cues, and is elevated in failing human left ventricle across distinct disease etiologies.

### LTBP2 Deficiency Improves Survival and Preserves Systolic Mechanics of the remote non-infarct segments of left ventricle after Myocardial Infarction

Although LTBP2 is required for microfibrillar development and stabilization in the eye (12), its role as a modifier of ECM remodeling in the heart remains poorly defined. Given its robust induction in activated CFs after MI, we asked whether global deletion of *Ltbp2* alters post-infarct remodeling and cardiac function. Wild-type (WT) and *Ltbp2* knockout (KO) mice were subjected to permanent left anterior descending coronary artery ligation and followed for 28 days. Kaplan–Meier analysis demonstrated significantly improved survival in KO mice compared with WT controls (Figure 2A).

**Figure 2.**
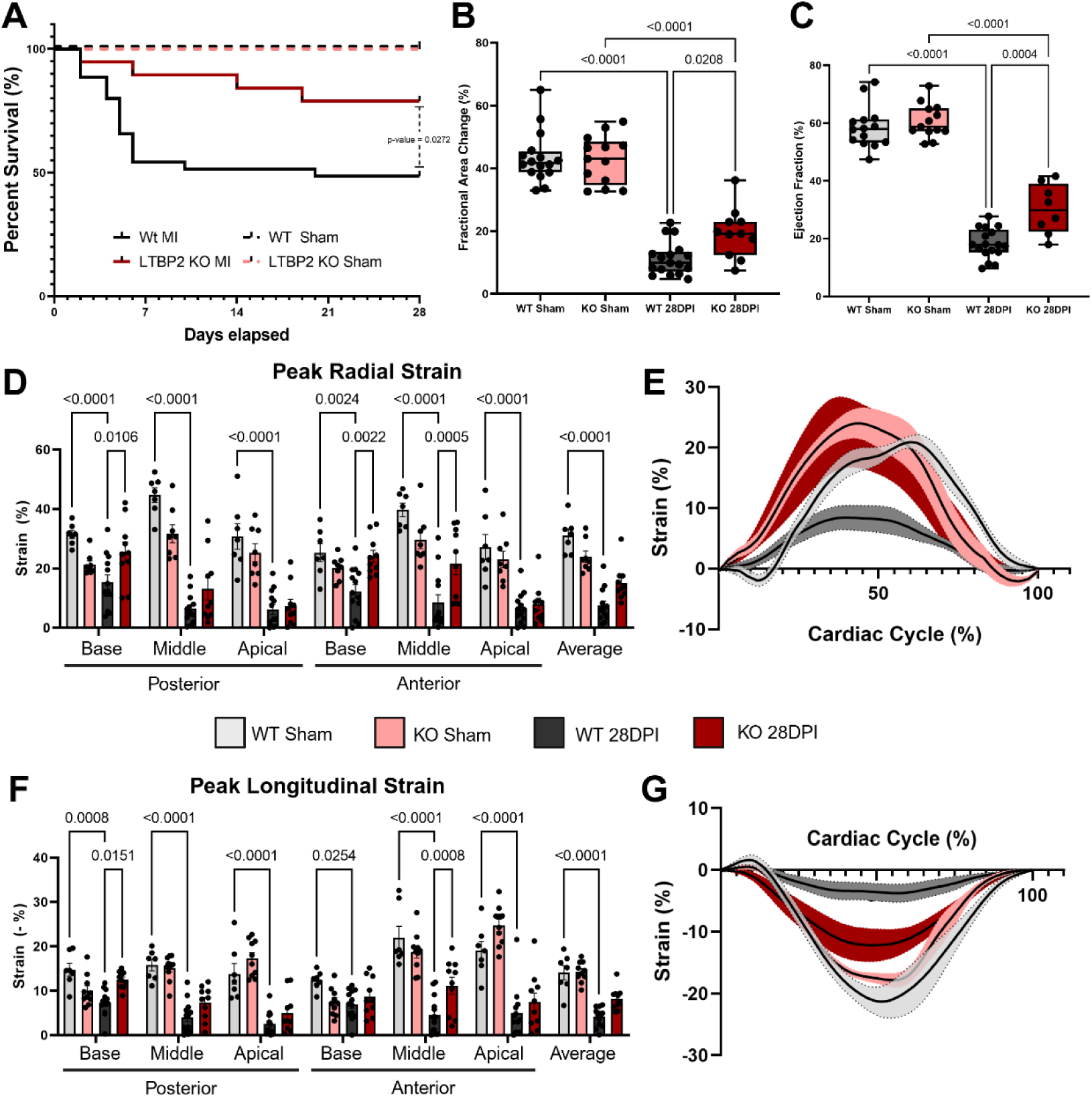
LTBP2 deficiency improves survival and preserves regional left ventricular mechanics 28 days after myocardial infarction. (A) Kaplan–Meier survival analysis of wild-type (WT) and *Ltbp2* knockout (KO) mice followed for 28 days after sham or myocardial infarction (MI) surgery (WT sham, n = 18; KO sham, n = 12; WT MI, n = 35; KO MI, n = 19). Survival was compared between WT MI and KO MI by log-rank (Mantel–Cox) test; P = 0.0272. (B) Fractional area change measured from the parasternal long-axis view using electrocardiogram-gated kilohertz visualization (EKV) at 28 days post-MI. N represented by each dot. (C) Left ventricular ejection fraction quantified by M-mode imaging at the mid-ventricular level at 28 days post-MI. N represented by each dot. (D) Regional endocardial peak radial strain assessed by two-dimensional speckle-tracking echocardiography (VevoStrain) from long-axis cines, reported for basal, mid, and apical segments of the posterior and anterior walls and as the segment-averaged value. (E) Mean radial strain across a single cardiac cycle for the anterior basal segment, plotted as a percentage of cycle duration. Solid lines show the group mean and shaded regions the SEM (n = 5-7 per group; WT MI versus KO MI peak strain, P = 0.0138). (F) Regional endocardial peak longitudinal strain quantified by VevoStrain from the same long-axis cines, plotted as absolute values; segments and averaging as in D. (G) Mean longitudinal strain across a single cardiac cycle for the anterior mid-ventricular segment (n = 5–7 per group; WT MI versus KO MI peak strain, P = 0.0179). Data in B and C are shown as box-and-whisker plots displaying the median, interquartile range, and full range, with individual animals overlaid; data in D and F are mean ± SEM with individual animals overlaid. For B and C, groups were compared by one-way ANOVA with Tukey post hoc testing. For D and F, comparisons were made by two-way ANOVA with genotype/surgical group and segment as factors, followed by Tukey post hoc testing. Only significant pairwise comparisons are annotated. All analyses were performed in GraphPad Prism.

To establish whether this survival benefit reflected differences in initial injury, we characterized both genotypes at the early post-infarct timepoint. Infarct size assessed by TTC staining was comparable between WT and KO hearts and both genotypes showed equivalent left ventricular dilation, with similar end-diastolic and end-systolic areas and a comparable, marked reduction in fractional area change and ejection fraction relative to sham controls at 7 days post MI (Supplementary Figure 2A–D). Regional wall thickness did not differ between genotypes at basal, mid, or apical levels (Supplementary Figure 2E), and speckle-tracking analysis showed equivalent depression of peak radial and longitudinal strain across all segments (Supplementary Figure 2F–G). WT and KO hearts did not differ significantly in both the extent of ischemic injury and the resulting early functional impairment, indicating that the divergence in survival was not attributable to differences in initial injury severity or gross hypertrophic remodeling.

By 28 days post-MI, however, KO mice demonstrated significant preservation of global systolic function, with improvements in both fractional area change and ejection fraction compared with WT MI hearts (Figure 2B–C). To determine whether these improvements were region-specific, we used two-dimensional speckle-tracking echocardiography to assess segmental myocardial mechanics. Radial strain analysis revealed significantly greater systolic strain in KO MI hearts at the middle and basal segments of the anterior wall and at the basal posterior wall, whereas apical segments corresponding to the infarcted territory remained similarly depressed in both genotypes (Figure 2D). Group-averaged radial strain curves across the cardiac cycle showed the same separation at the anterior basal segment (Figure 2E). Longitudinal strain followed the identical regional pattern, with improved deformation at posterior basal and anterior mid-wall segments in KO MI hearts (Figure 2F–G).

The regional distribution of this protection is notable. Improvement was confined to segments remote from the apical infarct, while the infarcted territory itself remained equally impaired in both genotypes, a pattern inconsistent with reduced ischemic injury and instead suggesting that LTBP2 acts on the surviving myocardium surrounding the scar. Together with the absence of any genotype-based difference at 7 days, these findings indicate that *Ltbp2* deletion improves post-MI survival and improved regional systolic function by limiting secondary remodeling of viable myocardium.

### LTBP2 Deficiency Enhances Remote Myocardial Diastolic Relaxation After Myocardial Infarction

The regional pattern of systolic protection in *Ltbp2*-deficient hearts, concentrated in remote, viable segments rather than in the infarct, suggested that LTBP2 acts on the mechanical properties of surviving myocardium. Because the passive properties of remote tissue are most directly interrogated during relaxation, and because chronic post-infarction remodeling is characterized by abnormal diastolic function as well as impaired contraction, we next examined whether deletion of *Ltbp2* alters regional diastolic mechanics.

Instantaneous strain and strain rate were exported across the cardiac cycle from the same speckle-tracking acquisitions at 28 days post-MI. Group-averaged radial strain curves showed reduced diastolic relaxation in WT MI hearts relative to sham controls, with a shallower descending limb indicating slower return toward baseline, whereas KO MI hearts retained relaxation kinetics closer to sham (Figure 3A–B). Quantification of peak diastolic radial strain rate confirmed significant impairment in WT MI hearts at the basal and mid-ventricular levels of both the posterior and anterior walls, with KO MI hearts showing significantly faster diastolic strain rates at all four of these segments (Figure 3C). The same pattern held in the longitudinal axis. WT MI hearts exhibited marked depression of the diastolic longitudinal strain rate, consistent with impaired relaxation, while KO MI hearts demonstrated significantly improved diastolic longitudinal strain rate at posterior and anterior basal and mid-ventricular segments (Figure 3D–F). Because these segments lie outside the infarcted apical territory, the improvement in relaxation is localized to viable myocardium, mirroring the regional distribution of the systolic findings and reinforcing that LTBP2 influences the mechanical behavior of remote tissue rather than the contractile performance of the scar.

**Figure 3.**
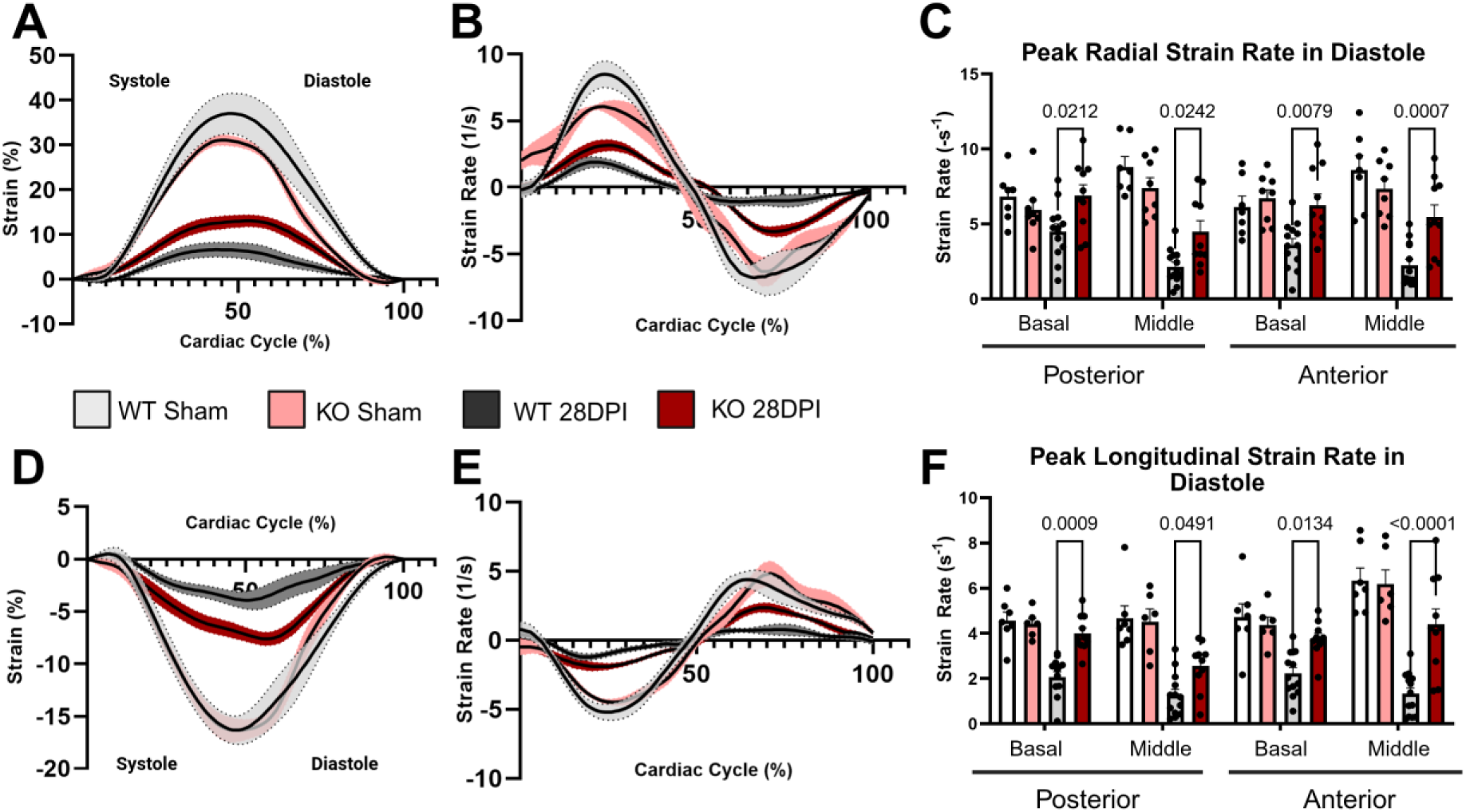
LTBP2 deficiency improves diastolic strain rate in the remote myocardium 28 days after myocardial infarction. (A) Mean radial strain across a single cardiac cycle, plotted as a percentage of cycle duration, with the vertical line marking end-systole and separating the systolic contraction and diastolic relaxation phases. Solid lines show the group mean and shaded regions the SEM. (B) Mean radial strain rate across the same cardiac cycle. Positive values correspond to systolic thickening and negative values to diastolic thinning; the peak negative deflection defines the peak diastolic radial strain rate quantified in C. (C) Peak diastolic radial strain rate in the posterior and anterior walls at basal and mid-ventricular levels segments remote from the apical infarct at 28 days after myocardial infarction (MI). (D) Mean longitudinal strain across a single cardiac cycle, presented as in A. (E) Mean longitudinal strain rate across the cardiac cycle, presented as in B; the peak positive deflection during diastole defines the peak diastolic longitudinal strain rate quantified in F. (F) Peak diastolic longitudinal strain rate in the posterior and anterior walls at basal and mid-ventricular levels at 28 days post-MI. Groups are wild-type (WT) sham, *Ltbp2* knockout (KO) sham, WT 28 days post-infarction (28DPI), and KO 28DPI, as indicated in the legend. Data in C and F are mean ± SEM with individual animals overlaid; n = 7-11 per group. Comparisons were made by two-way ANOVA with group and segment as factors, followed by Tukey post hoc multiple comparisons testing; only significant comparisons are annotated. All analyses were performed in GraphPad Prism.

### LTBP2 promotes remote interstitial fibrosis and tissue stiffening without altering cardiomyocyte hypertrophy

The functional data predicted a structural correlate: if *Ltbp2* deletion improves relaxation of viable myocardium without altering infarct size, the difference should reside in the remote interstitium. Picrosirius Red staining of remote myocardium at 28 days post-MI revealed extensive interstitial collagen deposition in WT hearts, distributed diffusely between cardiomyocytes rather than as a confluent scar, consistent with reactive fibrosis of viable tissue. KO hearts showed markedly less collagen, with percent fibrotic area not significantly different from sham (Figure 4A–B). Remote remodeling after infarction involves cardiomyocyte hypertrophy as well as matrix expansion. Wheat germ agglutinin staining of remote myocardium showed increased cardiomyocyte cross-sectional area in both WT and KO hearts at 28 days post-MI relative to their respective shams, with no difference between genotypes (Figure 4C–D). The hypertrophic response of surviving cardiomyocytes is therefore intact in *Ltbp2*-deficient hearts, localizing the effect of LTBP2 to the interstitial compartment rather than to the myocyte. Because histology reports how much matrix is present rather than how the tissue behaves mechanically, we next measured regional stiffness directly by atomic force microscopy–based nanoindentation on vibratome-cut sections. WT MI hearts showed substantially increased tissue stiffness in both regions, with infarct exceeding remote, consistent with scar maturation superimposed on diffuse remote remodeling. *Ltbp2*-deficient hearts were significantly less stiff in both regions, with KO remote tissue approaching sham levels (Figure 4E). Sham stiffness and sham fibrosis did not differ between genotypes, indicating that LTBP2 acts on injury-induced remodeling rather than baseline matrix mechanics. Because cardiomyocyte size was matched between genotypes, the difference in tissue stiffness in the remote region is likely attributable to the composition of the interstitium rather than to a difference in the cellular makeup of the tissue being indented. Reduced collagen and preserved compliance in remote myocardium provide a direct physical basis for the faster diastolic strain rates and better-preserved regional deformation in KO hearts, and the preserved hypertrophic response argues against a generalized attenuation of remodeling in *Ltbp2*-deficient animals. However, these measurements report the endpoint of fibrotic remodeling without identifying the programs that produce it.

**Figure 4.**
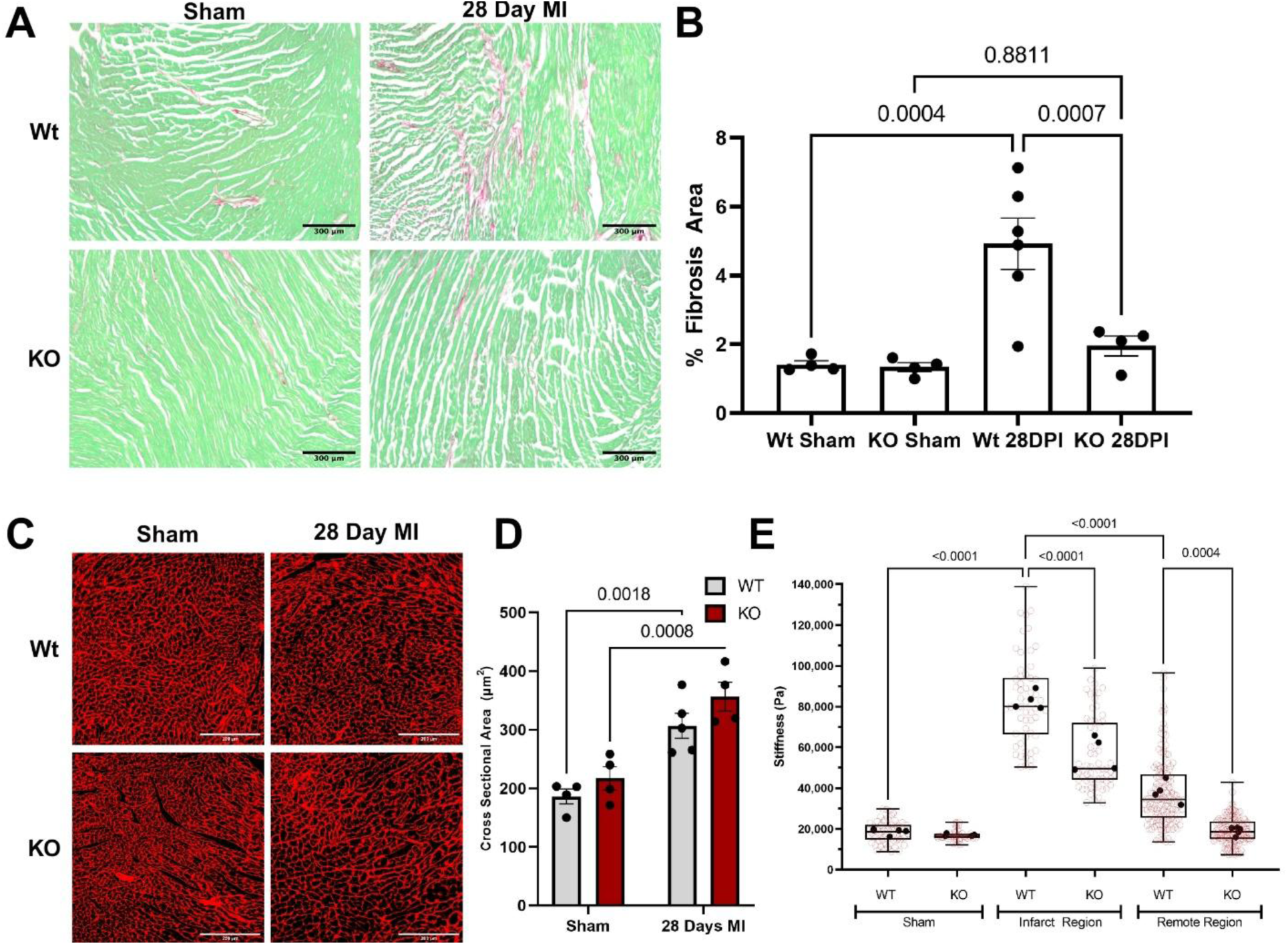
LTBP2 deficiency attenuates remote myocardial fibrosis and reduces tissue stiffness 28 days after myocardial infarction. (A) Representative Picrosirius Red/Fast Green staining of remote myocardium from wild-type (WT) and *Ltbp2* knockout (KO) hearts at 28 days after sham surgery or myocardial infarction (MI). Collagen is stained red and myocardium green. Scale bars, 150 μm. (B) Quantification of percent fibrotic area in the remote region. Image acquisition and quantification were performed by an investigator blinded to genotype and surgical group. Data are mean ± SEM with individual animals overlaid; n = 4–6 per group. (C) Wheat germ agglutinin (WGA) staining of cardiac sections at 28 days post MI. (D) Cardiomyocyte cross sectional area measurement taken from WGA staining. (E) Atomic force microscopy measurements of myocardial elastic modulus (kPa) in infarct and remote regions of WT and KO hearts at 28 days post-MI, with sham hearts as controls. Black filled points represent per-animal means (biological replicates, n = 4–5 per group); light red open points represent individual force–indentation curves acquired across repeated force maps (technical replicates). Boxes show the median and interquartile range with whiskers spanning the full range. Testing was performed with the operator blinded to both genotype and tissue region, and WT and KO samples were interleaved across sessions. Comparisons were made by one-way ANOVA with Tukey post hoc multiple comparisons testing; only annotated comparisons are shown. All analyses were performed in GraphPad Prism.

### LTBP2 Promotes Activation of TGF-β Associated Fibrotic Signaling in Remote Myocardium After MI

To define the molecular programs driving remodeling within remote myocardium, we performed unbiased proteomic profiling of remote left ventricular tissue from WT and KO hearts at 28 days post-MI, with sham-operated hearts as controls. Principal component analysis separated the four groups, with WT MI forming the most distinct cluster and KO MI positioned closer to the sham groups, indicating that loss of *Ltbp2* attenuates the magnitude of the proteomic remodeling in the remote myocardium (Figure 5A). Unsupervised clustering of all quantified proteins showed the same structure (Supplementary Figure 3A).

**Figure 5.**
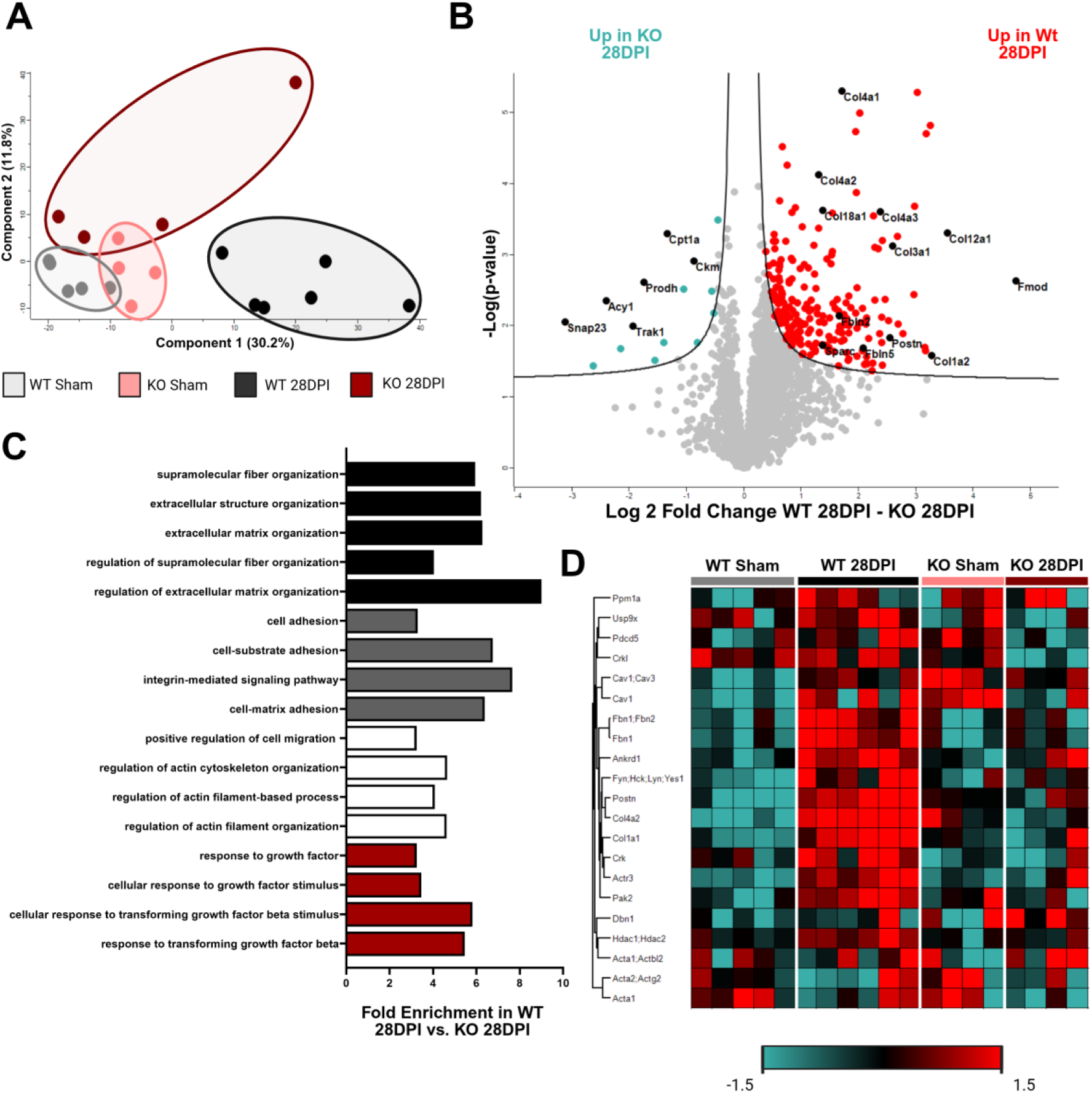
Proteomic profiling of remote myocardium identifies enhanced TGF-β–associated fibrotic signaling in wild-type hearts 28 days after myocardial infarction. (A) Principal component analysis of label-free quantitative proteomes from remote left ventricular myocardium of wild-type (WT) and *Ltbp2* knockout (KO) mice at 28 days after sham surgery or myocardial infarction (MI). The first two components account for 30.2% and 11.8% of total variance. Each point represents one animal; n = 4-6 per group. (B) Volcano plot of differentially abundant proteins in WT versus KO remote myocardium at 28 days post-MI, plotting log₂ fold change against −log₁₀ P value. Red points denote proteins significantly increased in WT and teal points those increased in KO; curved lines indicate the significance boundary. Selected extracellular matrix and fibrosis-associated proteins are labelled. Significance was determined by two-tailed Student’s t-test with permutation-based false discovery rate correction (FDR < 0.05, s0 = 0.01) in Perseus. (C) Gene ontology biological process enrichment analysis of proteins significantly increased in WT relative to KO remote myocardium, ranked by fold enrichment. All biological processes had Benjamin-Hochberg adjusted p-value of less than 0.05. Analysis performed using PANTHER against full mouse proteome. (D) Heat map of proteins annotated under the gene ontology term “cellular response to transforming growth factor beta stimulus,” with rows clustered by Euclidean distance and columns grouped by experimental condition. Each column represents an individual animal. Colour reflects z-scored protein intensity across samples, from −1.5 (teal) to 1.5 (red).

WT MI remote myocardium was significantly enriched for fibrosis- and ECM-associated proteins relative to WT sham, including fibrillar Collagens, Periostin, Fibromodulin, Fibrillin-1, and microfibril-associated proteins, confirming that the remote, viable myocardium, undergoes substantial chronic matrix remodeling beyond just the acute infarct healing period (Supplementary Figure 3B). The equivalent comparison in KO hearts yielded no proteins meeting the FDR < 0.05 threshold, when compared to sham KO controls, indicating that *Ltbp2* deficiency largely suppresses this chronic remodeling program (Supplementary Figure 3C). Direct comparison of WT MI and KO MI remote myocardium confirmed enrichment in WT hearts of fibrillar collagens Col1a1, Col1a2, Col3a1, and Col12a1, basement membrane collagens Col4a1–3, matricellular proteins Postn and Sparc, and microfibril-associated regulators including Fbln2 and Fbln5 (Figure 5B). Gene ontology analysis of these proteins identified extracellular matrix organization, elastic fiber assembly, and integrin-mediated signaling among the most enriched biological processes and, notably, cellular response to transforming growth factor-β stimulus (Figure 5C). Hierarchical clustering of proteins annotated under that term showed coordinated upregulation across WT MI samples relative to KO MI and both sham groups (Figure 5D). TGF-β pathway engagement therefore emerges as a distinguishing feature of the WT remote remodeling response and offers a candidate explanation for the divergence in fibrosis, stiffness, and mechanical function between genotypes. It also presents a mechanistic problem.

### LTBP2 Enhances TGF-β Bioavailability by Limiting Extracellular Matrix Sequestration

To resolve how an LTBP that does not bind latent TGF-β amplifies TGF-β signaling, we turned to primary CFs. Published single-cell RNA sequencing data showed that *Ltbp2* and *Fbn1* peak during the early activated fibroblast phase after injury, whereas *Ltbp1* is highest in healthy myocardium and rises again in later-stage fibroblast states (Figure 6A). RT-qPCR of cultured CFs reproduced this temporal ordering, with *Ltbp2* and *Fbn1* induced by day 7 and *Ltbp1* rising later (Figure 6C), indicating that matrix maturation in culture recapitulates the sequence of microfibrillar gene expression seen in vivo.

**Figure 6.**
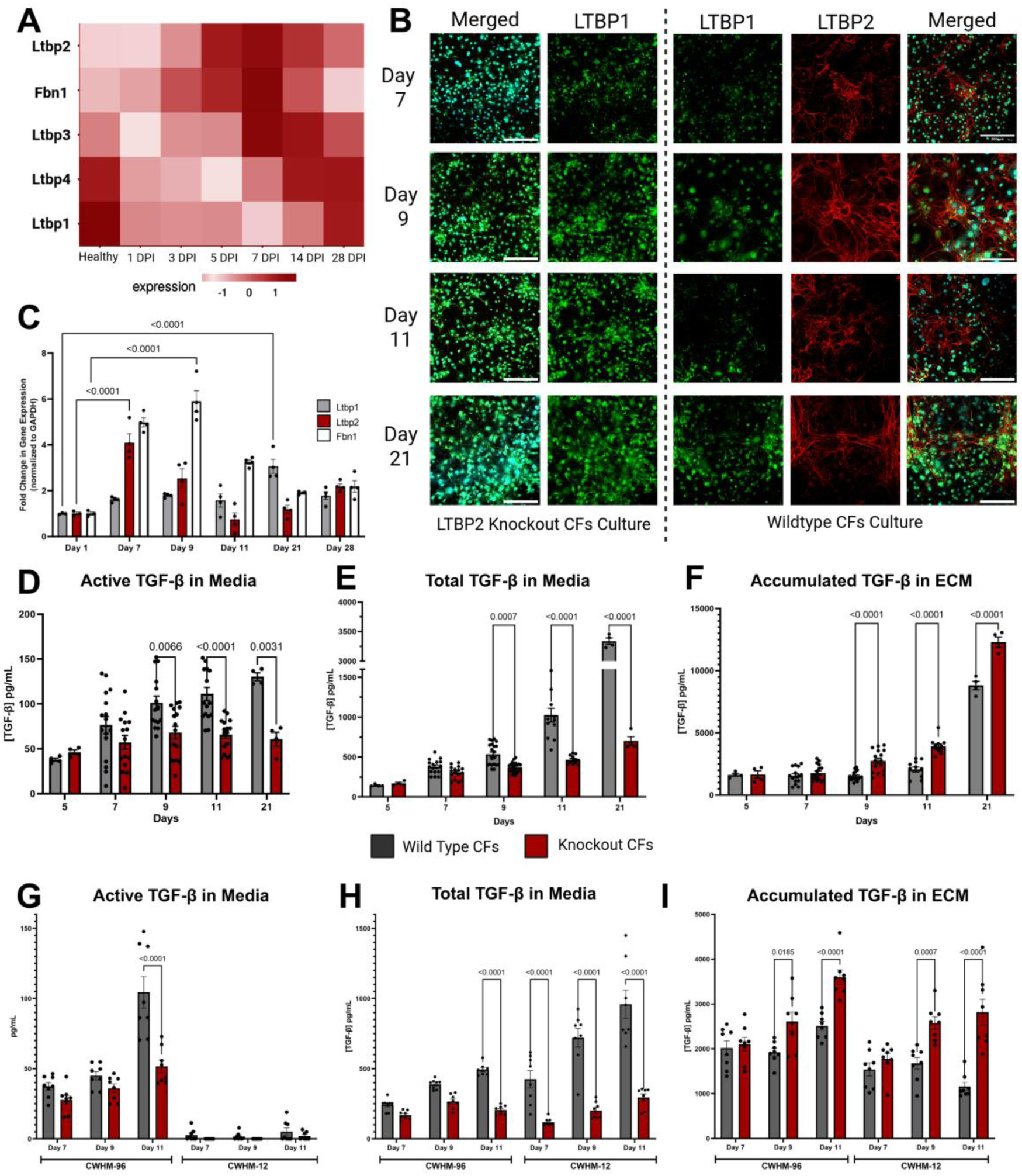
LTBP2 limits extracellular matrix sequestration of latent TGF-β and increases TGF-β bioavailability in cultured cardiac fibroblasts. (A) Expression of Ltbp family genes and Fbn1 in cardiac fibroblasts from healthy hearts and at 1, 3, 5, 7, 14, and 28 days post-infarction (DPI), derived from published single-cell RNA sequencing data (Patrick et al. 2025) (48). Colour represents scaled mean expression per timepoint. (B) Representative immunofluorescence of Ltbp2 knockout (KO, left) and wild-type (WT, right) primary cardiac fibroblast cultures at days 7, 9, 11, and 21 after plating, stained for LTBP1 (green), LTBP2 (red), and nuclei (DAPI, cyan). Images are representative of n > 4 independent isolations. Scale bars, 200 μm. (C) RT-qPCR of Ltbp1, Ltbp2, and Fbn1 in WT cardiac fibroblast cultures at days 1, 7, 9, 11, 21, and 28 after plating, normalized to Gapdh and expressed as fold change relative to day 1. n = 4 independent isolations per timepoint. (D–F) Conditioned media and cell-derived matrix were collected from WT and KO cardiac fibroblast cultures at the indicated days after plating and assayed for TGF-β activity using the transformed mink lung epithelial cell (TMLC) PAI-1 promoter–luciferase reporter. (D) Active TGF-β in conditioned media, measured directly. (E) Total TGF-β in conditioned media, measured after heat activation (80°C, 10 min) to convert latent to active ligand. (F) Matrix-associated TGF-β, released from decellularized matrix by acid activation and neutralization. (G–I) Wild-type and Ltbp2 knockout primary cardiac fibroblasts were treated with the pan–αv integrin inhibitor CWHM-12 (5 μM) or its inactive enantiomer CWHM-96 (vehicle control) for 24 hours before collection at days 7, 9, and 11 after plating. TGF-β was quantified by TMLC reporter assay as in D–F. (G) Active TGF-β measured directly from conditioned media. (H) Total TGF-β in conditioned media after heat activation (80°C, 10 min). (I) Matrix-associated TGF-β released from decellularized ECM by acid activation and neutralization. Data are mean ± SEM with individual replicates overlaid. For C–F, comparisons were made by two-way ANOVA followed by Tukey post hoc testing; for G–I, by one-way ANOVA with Tukey post hoc testing. Only significant comparisons are annotated. All analyses were performed in GraphPad Prism.

Immunofluorescence over time showed progressive deposition of LTBP2 into fibrillar structures in WT cultures, with comparatively sparse LTBP1 incorporation, whereas *Ltbp2*-deficient cultures assembled visibly more extensive LTBP1 networks (Figure 6B). Skeleton analysis quantified this difference: by day 21, both average branch length and total skeleton length of LTBP1-positive structures were significantly greater in KO than WT cultures (Supplementary Figure 4A–B). To test whether this altered composition changes TGF-β handling, TGF-β was measured in conditioned media and matrix fractions by TMLC reporter assay. WT CFs cultures contained significantly more bioactive TGF-β than KO CFs from day 9 onward (Figure 6D) and more total TGF-β at later timepoints, with a pronounced increase at day 21 coinciding with the rise in *Ltbp1* expression (Figure 6E). KO CFs showed the reciprocal pattern, accumulating significantly more TGF-β within the matrix at the same timepoints (Figure 6F). In the absence of LTBP2, latent TGF-β is therefore retained in the matrix rather than released into the soluble compartment.

To determine how the released ligand becomes active, we inhibited αv integrins with CWHM-12, using its inactive enantiomer CWHM-96 as control. Integrin blockade reduced active TGF-β to near-undetectable levels in both genotypes (Figure 6G), confirming that activation is integrin-dependent. The genotype difference in compartmentalization nonetheless persisted, with WT cultures retaining greater soluble and lower matrix-associated TGF-β than KO cultures under inhibition (Figure 6H–I). LTBP2-dependent redistribution therefore operates upstream of, or in parallel with, integrin-mediated activation rather than requiring it.

### The LTBP2 fibrillin-1–binding module binds FBN1 directly and is sufficient to mobilize matrix-sequestered TGF-β

The preceding data establish that loss of LTBP2 retains latent TGF-β in the matrix, but they do not establish sufficiency, nor do they confirm that LTBP2 engages fibrillin-1 in this system. To address both, we generated a construct spanning EGF13–16 of human LTBP2, corresponding to the fibrillin-1–binding module previously mapped by Dr. Nakmura’s Lab (15), and delivered it either as purified protein or by lentiviral expression (Figure 7A). The fragment used here is shorter than the previously reported binding region, and in a cell-free assay the purified EGF13–16 module was sufficient to co-precipitate human FBN1 rF23, confirming that this minimal module retains direct fibrillin-1 binding (Figure 7B).

**Figure 7.**
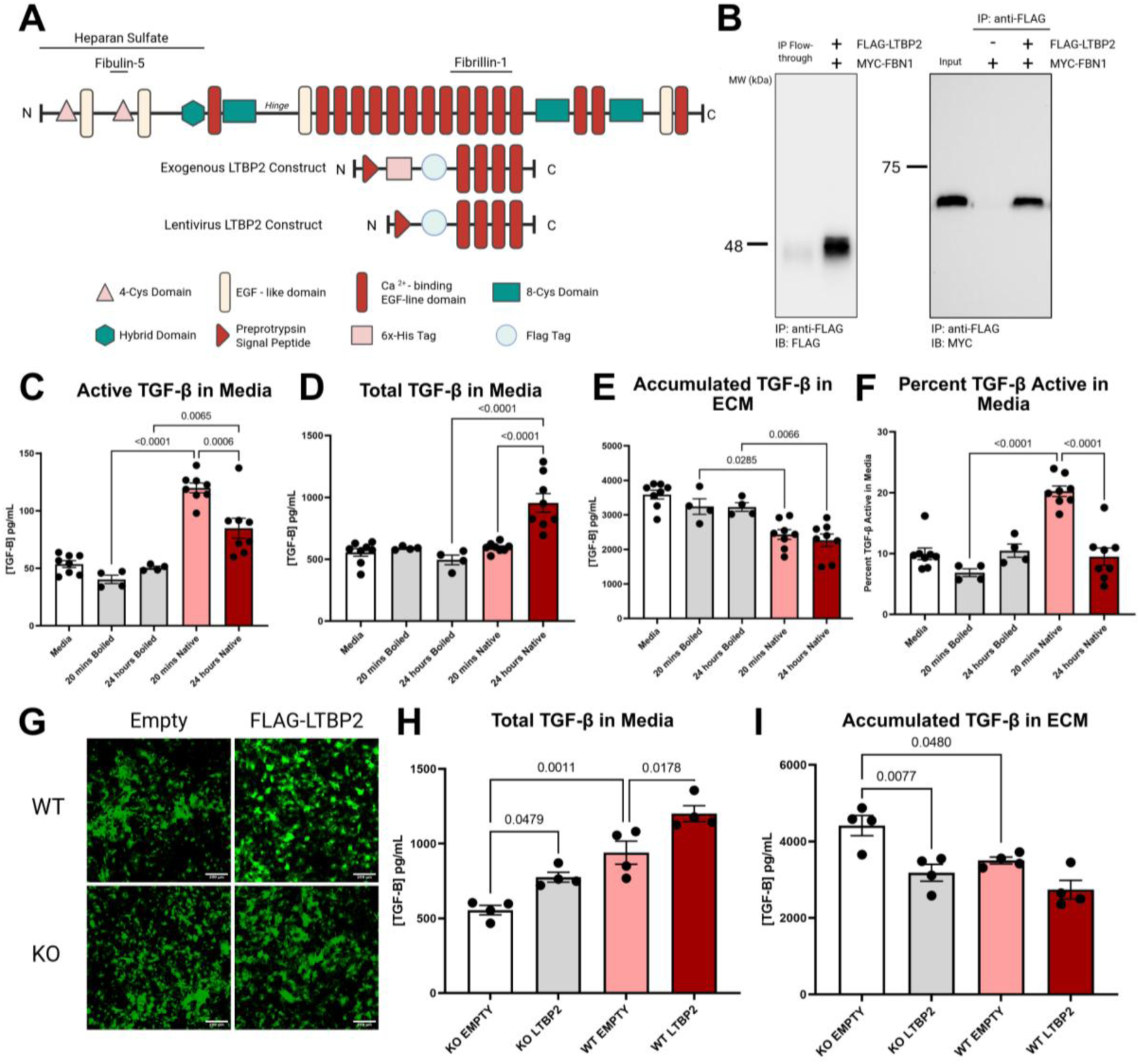
The LTBP2 EGF13–16 fibrillin-1–binding module binds FBN1 directly and is sufficient to shift TGF-β from the matrix to the soluble compartment in cardiac fibroblasts. (A) Domain organization of full-length LTBP2 showing previously mapped binding regions for heparan sulfate, fibulin-5, and fibrillin-1, and the two EGF13–16 constructs used here. The exogenous construct carries an N-terminal preprotrypsin signal peptide, 6×His and FLAG tags; the lentiviral construct is identical except that the His tag is omitted. Both span a shorter region than the previously reported fibrillin-1–binding fragment (15). (B) Cell-free co-immunoprecipitation of purified FLAG-tagged LTBP2 EGF13–16 with purified MYC-tagged FBN1 rF23 (15). Proteins were combined in calcium-containing buffer, captured on anti-FLAG beads, and immunoblotted for FLAG (left) or MYC (right). Left: FLAG-LTBP2 partitions into the bound fraction relative to the IP flow-through. Right: MYC-FBN1 is recovered only when FLAG-LTBP2 is present, with input shown for reference. (C–F) Purified LTBP2 EGF13–16 (1.5 μM) or heat-denatured protein was added to Ltbp2-knockout primary cardiac fibroblast cultures at day 7 after plating, for 20 minutes or 24 hours; medium alone served as an additional control. TGF-β was quantified using the TMLC PAI-1 promoter–luciferase reporter. (C) Active TGF-β in conditioned medium. (D) Total TGF-β in conditioned medium, measured after heat activation. (E) Matrix-associated TGF-β released from decellularized ECM by acid activation. (F) Proportion of total medium TGF-β in the active form. Native protein increased active and total soluble TGF-β and reduced matrix-associated TGF-β, whereas heat-denatured protein was indistinguishable from medium alone at both timepoints. n = 4–8 independent cultures per condition. (G) Representative GFP fluorescence in wild-type (WT) and Ltbp2-knockout (KO) primary cardiac fibroblasts transduced with empty or FLAG-LTBP2 EGF13–16 lentivirus, confirming comparable transduction across groups. Scale bars, 200 μm. (H, I) TGF-β distribution in WT and KO cardiac fibroblasts at 11 days after plating, 4 days after transduction. (H) Total TGF-β in conditioned medium released through heat activation (80°C, 10mins). (I) Matrix-associated TGF-β. Expression of LTBP2 EGF13–16 increased soluble and decreased matrix-associated TGF-β in both genotypes. Data are mean ± SEM with individual replicates overlaid. Comparisons in C–F were made by one-way ANOVA with Tukey post hoc testing; in H–I by two-way ANOVA with genotype and construct as factors, followed by Tukey post hoc testing. Only significant comparisons are annotated. All analyses were performed in GraphPad Prism.

Addition of purified EGF13–16 to *Ltbp2*-null CFs reproduced the wild-type TGF-β distribution. Active TGF-β in conditioned media rose sharply within 20 minutes and remained elevated at 24 hours (Figure 7C), while matrix-associated TGF-β fell at both timepoints (Figure 7E) and the proportion of media TGF-β in the active form doubled acutely (Figure 7F). Heat-denatured protein was indistinguishable from medium alone across all readouts, confirming that the effect requires the folded module. The kinetics are informative: the immediate rise in active ligand at 20 minutes preceded any increase in total soluble TGF-β, which appeared only by 24 hours (Figure 7D). Lentiviral expression of the same module tested whether this held when LTBP2 is produced by the cell during matrix assembly. Transduction efficiency was comparable across genotypes and constructs (Figure 7G). Expression of EGF13–16 increased total TGF-β in conditioned media and decreased matrix-associated TGF-β in both wild-type and *Ltbp2*-null fibroblasts (Figure 7H–I), with empty-vector controls reproducing the genotype difference seen in Figure 6. Notably, the construct shifted TGF-β distribution in wild-type cells that already possess an intact microfibrillar system, indicating that LTBP2 abundance is rate-limiting for matrix sequestration rather than merely permissive. Consistent with this, proteomic profiling of cell-derived matrix showed that wild-type fibroblasts deposit a progressively more profibrotic matrix than knockouts, with enrichment of FBN1, FBN2, MFAP2, THBS2, and fibrillar collagens by day 21 (Supplementary Figure 5).

Together with the loss-of-function data, these results complete the mechanism. LTBP2 binds fibrillin-1 through a minimal EGF13–16 module, and introducing that module alone, acutely as purified protein or by sustained expression, is sufficient to shift latent TGF-β from the matrix into the soluble, activatable compartment. This provides a molecular explanation for the TGF-β pathway enrichment, interstitial fibrosis, myocardial stiffening, and functional decline observed in wild-type remote myocardium after MI and resolves how an LTBP lacking latent TGF-β–binding capacity nonetheless amplifies TGF-β signaling.

## Discussion

In this study, we identify LTBP2 as a previously unrecognized mediator of maladaptive remodeling in the remote myocardium following MI. While post-infarction remodeling has traditionally focused on the infarct and peri-infarct regions, our findings highlight the importance of molecular and structural changes that extend beyond the primary site of ischemic injury. We demonstrate that injury-induced LTBP2 expression in activated cardiac fibroblasts promotes ECM remodeling, myocardial stiffening, and functional deterioration in viable myocardium distant from the infarct scar. Genetic deletion of *Ltbp2* improved survival, preserved systolic and diastolic mechanics, reduced fibrosis and myocardial stiffness, and attenuated TGF-β–associated profibrotic signaling networks in the remote myocardium.

Mechanistically, we show that LTBP2 binds fibrillin-1 through a minimal EGF13–16 module and that introducing this module alone is sufficient to mobilize latent TGF-β out of the matrix and into the soluble, activatable compartment, establishing both necessity and sufficiency for a protein that cannot itself bind latent TGF-β. LTBP2 is likewise elevated in failing human left ventricle across ischemic, non-ischemic, and hypertrophic cardiomyopathy, indicating that this pathway is engaged in human disease irrespective of the initiating insult.

### LTBP2 promotes remote interstitial remodeling and tissue stiffening after MI

Our data reveal that LTBP2 is rapidly induced following MI and deposited throughout both infarct and remote myocardial regions. Although scar formation within the infarct is an expected component of wound healing, remodeling of the remote myocardium is a major determinant of long-term ventricular function and progression to heart failure. Remote myocardial fibrosis contributes directly to increased ventricular stiffness, impaired relaxation, and progressive ventricular dilation, even in regions not directly exposed to ischemic injury (33, 34). Consistent with this paradigm, we observed robust induction of LTBP2 in injury-activated fibroblasts, accompanied by deposition within fibrillin-1–rich ECM structures in the remote myocardium.

Genetic deletion of *Ltbp2* did not alter infarct size, ventricular dilation, wall thickness, or regional strain at 7 days post-MI, establishing that the two genotypes sustained equivalent ischemic injury and equivalent early functional impairment. The divergence in survival, fibrosis, stiffness, and mechanics emerged only over the subsequent three weeks, following peak LTBP2 expression at 7 days, indicating that LTBP2 contributes to the secondary remodeling processes rather than the acute response to ischemia.

A key observation is where that divergence occurs, and where it does not. Cardiomyocyte cross-sectional area in remote myocardium increased comparably in both genotypes after MI, so the hypertrophic response of surviving myocytes to altered loading is fully intact in *Ltbp2*-deficient hearts. This dissociation is informative in two respects. It argues against a model in which knockout hearts simply have less global remodelling due to milder initial injury, both genotypes mount the same load-driven cellular hypertrophy and differ only in the interstitium through the extent of ECM deposition. It also localizes the effect of LTBP2 to the matrix compartment specifically, which is precisely what a microfibril-centered mechanism predicts, and it suggests the reduced elastic modulus we measured by atomic force microscopy reflects interstitial composition rather than a different cellular makeup of the indented tissue.

Knockdown of *Ltbp2* has previously been reported to attenuate cardiac fibrosis in dilated cardiomyopathy and angiotensin II–induced hypertrophy models, where the effects were attributed to NF-κB signaling and oxidative stress (35, 36). Our findings extend these observations to ischemic injury and to the remote, non-infarcted compartment specifically, and identify a distinct upstream mechanism centered on microfibrillar composition and TGF-β compartmentalization. Whether the NF-κB and oxidative stress signatures described in those models represent parallel pathways or downstream consequences of altered TGF-β bioavailability remains to be determined.

The relationship between fibrosis and stiffness is likely bidirectional in this setting. Increased ECM accumulation and collagen remodeling directly increase myocardial stiffness, which impairs ventricular filling and alters mechanical loading. In turn, increased tissue stiffness activates fibroblasts through mechanotransductive pathways, reinforcing ECM deposition and sustaining profibrotic signaling (37). Our own data place LTBP2 within this loop: *Ltbp2* was itself induced by increased substrate stiffness in primary CFs, positioning it as both a product and a driver of the stiffening response. LTBP2 may therefore contribute to a self-amplifying cycle in which fibroblast activation promotes matrix deposition, matrix deposition increases stiffness, and increased stiffness further induces LTBP2 and sustains pathologic remodeling.

Together, these findings implicate LTBP2 not merely as a marker of injury-activated fibroblasts, but as a mechanistic driver of fibrotic remodeling at sites distant from the ischemic core.

### LTBP2 regulates TGF-β bioavailability through the extracellular matrix

TGF-β is a central regulator of fibroblast activation, ECM deposition, and post-injury remodeling, and fibroblast-specific deletion of TGF-β receptors or Smad3 is sufficient to attenuate cardiac fibrosis, establishing fibroblast-autonomous TGF-β signaling as a principal driver of the fibrotic response (37). During biosynthesis, TGF-β is secreted as a small latent complex in which mature TGF-β remains noncovalently bound to its latency-associated peptide. This complex is covalently linked to a latent TGF-β–binding protein to form the large latent complex (LLC), which is subsequently deposited into the ECM. By tethering the LLC to fibrillin-rich microfibrils, LTBPs impose spatial and mechanical control over growth-factor activation, enabling context-dependent release of active TGF-β in response to integrin-mediated traction, proteolytic cleavage, or other matrix-dependent mechanisms (30).

Four LTBP isoforms are encoded by distinct genes and exhibit non-redundant matrix-binding properties and tissue distributions. LTBP1, LTBP3, and LTBP4 each covalently bind latent TGF-β: LTBP1 has been linked to vascular development and aneurysm susceptibility (38), LTBP3 to skeletal and craniofacial morphogenesis (39), and LTBP4 to elastogenesis and pulmonary homeostasis (40, 41). LTBP2 is unique within the family in that it does not covalently bind latent TGF-β and has instead been assigned structural roles in microfibril organization, elastogenesis, and integrin signaling. How LTBP2 nonetheless promotes TGF-β–dependent fibrosis, as we and others observe, has remained unresolved.

Our data support a model in which LTBP2 regulates TGF-β signaling indirectly, by remodeling the competitive binding environment of the microfibril. Several lines of evidence converge on this mechanism. LTBP family members and fibulins are known to compete for defined, site-specific binding positions on fibrillin-1, and LTBP2 has been shown biochemically to compete with LTBP1 for the fibrillin-1 N-terminal region (19, 42). Reducing the microfibrillar incorporation of LTBP1 is in turn sufficient to increase TGF-β bioavailability and signaling without altering total TGF-β expression, establishing that displacement of LTBP1-bound LLCs from microfibrils, rather than a change in ligand abundance, governs local TGF-β release (18, 43). Against this background, we find that loss of LTBP2 in cardiac fibroblasts produces exactly the reciprocal change in microfibrillar composition that competition predicts: *Ltbp2*-null cultures assembled significantly more extensive LTBP1-positive fibrillar networks, with greater branch length and total skeleton length by day 21, and the matrix they deposited differed compositionally from wild-type, which accumulated a progressively more profibrotic proteome including fibrillin-1, fibrillin-2, MFAP2, thrombospondin-2, and fibrillar collagens.

Two advances place this model on firmer ground than previously possible. First, we show that a minimal module spanning EGF13–16 of LTBP2, shorter than the fibrillin-1–binding region previously mapped (12), is sufficient to bind fibrillin-1 directly in a cell-free assay. This narrows the interaction to a defined domained and provides a site to potentially regulate its interaction. Second, and more consequentially, introducing that module alone into *Ltbp2*-null fibroblasts was sufficient to reproduce the wild-type TGF-β distribution. The kinetics of that response are themselves informative. Active TGF-β in conditioned media rose sharply within twenty minutes of adding the purified module, and the proportion of soluble ligand in the active form doubled over the same interval, while total soluble TGF-β did not increase until twenty-four hours. The immediate appearance of active ligand without a corresponding rise in the soluble pool suggests LTBP2 appears to remove latent complexes off the existing microfibrillar reservoir, making them immediately available for activation. Sustained expression of the same module by lentiviral delivery reproduced this redistribution, increasing soluble and decreasing matrix-associated TGF-β. Critically, it did so not only in knockout fibroblasts but in wild-type cells with a fully intact microfibrillar system. That raising LTBP2 above endogenous levels further displaces TGF-β from the matrix indicates that LTBP2 abundance is rate-limiting for matrix sequestration rather than merely permissive.

The latent complexes inhibited by LTBP2 remain dependent on downstream activation. αv-integrin blockade with CWHM-12 reduced active TGF-β to near-undetectable levels in both genotypes, yet the genotype difference in media-versus-matrix partitioning persisted. LTBP2-dependent compartmentalization therefore operates upstream of, or in parallel with, integrin-mediated activation rather than through it, and the two steps are mechanistically separable: LTBP2 determines how much latent ligand is available, integrins determine how much of it becomes active.

This mechanism offers a resolution to the long-standing paradox of how a non–TGF-β–binding LTBP amplifies TGF-β signaling. By competitively remodeling microfibril occupancy, injury-induced LTBP2 shifts the equilibrium from matrix sequestration toward sustained TGF-β bioavailability, reinforcing fibroblast activation, ECM deposition, and stiffening in a self-amplifying loop. In the remote myocardium, where fibroblasts are activated but injury is not driven by ischemic necrosis, this LTBP2-dependent shift may be a principal determinant of the maladaptive, TGF-β–driven fibrosis that distinguishes viable remote tissue from the infarct scar.

### Remote myocardial remodeling as a determinant of cardiac dysfunction

Our findings emphasize that remote myocardial remodeling is not merely a secondary consequence of infarct expansion, but an active pathologic process that contributes to post-MI functional decline. While infarct size remains an important predictor of early outcomes, progressive fibrosis and stiffening of remote myocardium are major contributors to long-term ventricular dysfunction and heart failure progression (1–3). Increased myocardial stiffness impairs ventricular filling, elevates diastolic pressures, alters wall stress, and promotes adverse ventricular remodeling.

The preservation of diastolic strain dynamics and myocardial compliance in *Ltbp2*-deficient hearts suggests that LTBP2 regulates the mechanical properties of post-MI myocardium. The concordance across independent modalities is what makes this argument strong: proteomic profiling identified ECM organization and TGF-β–responsive programs as the dominant differences between genotypes, histology showed reduced interstitial collagen, atomic force microscopy showed reduced elastic modulus, and speckle-tracking showed faster diastolic strain rates, with all four measures localizing the effect to the same remote, viable segments, none of them differing between genotypes at baseline or at 7 days, and cardiomyocyte hypertrophy explicitly unaffected.

An important nuance is that the remote proteome of *Ltbp2*-deficient hearts remained close to its own sham baseline after MI, whereas wild-type remote myocardium diverged markedly. Rather than shifting remodeling toward a different profibrotic program, loss of LTBP2 appears to blunt the magnitude of the response. This is consistent with a mechanism acting on the availability of an upstream driver, TGF-β, rather than on any single downstream effector, and with the in vitro finding that LTBP2 levels set the position of a sequestration equilibrium rather than switching a pathway on or off.

### Clinical relevance and therapeutic implications

The identification of LTBP2 as a regulator of remote myocardial remodeling has implications for understanding progression from myocardial injury to heart failure. Current therapeutic strategies after MI focus primarily on limiting infarct size, reducing ischemic injury, and suppressing neurohormonal activation (44). Few interventions directly target fibrotic remodeling of viable remote myocardium, despite its contribution to diastolic dysfunction, ventricular stiffening, and progressive heart failure.

Targeting LTBP2 may represent a strategy to attenuate pathologic fibrosis while avoiding global suppression of TGF-β signaling. This distinction matters because TGF-β is essential for tissue repair, immune regulation, and homeostasis (45), and systemic TGF-β inhibition has been limited by on-target toxicity. Rather than blocking the ligand or its receptor, modulating LTBP2 would act on where latent TGF-β resides within the matrix, restoring sequestration rather than eliminating signaling. In principle this preserves the spatially and mechanically gated activation that normal repair depends on. Two features of our data support the feasibility of such an approach. *Ltbp2*-null mice showed no baseline cardiac phenotype in function, fibrosis, cardiomyocyte size, or tissue stiffness, indicating that LTBP2 is largely dispensable in the uninjured heart and that a therapeutic window may exist. And because the relevant activity resides in a defined, minimal fibrillin-1–binding module, the interaction surface is small enough to be a plausible target for peptide or small-molecule interference rather than requiring suppression of an entire protein.

Our human data strengthen the case for relevance. LTBP2 abundance was significantly increased in left ventricular tissue from patients with ischemic, non-ischemic, and hypertrophic cardiomyopathy relative to non-failing controls, and immunofluorescence showed LTBP2 deposition overlapping FBN1-rich microfibrillar structures in diseased myocardium. That LTBP2 is elevated across etiologies with distinct primary insults, ischemic injury, primary myocardial disease, and pressure-driven hypertrophy, suggests it marks a convergent fibrotic response rather than an ischemia-specific one. Consistent with this, elevated LTBP2 has been reported in pulmonary fibrosis and as a plasma biomarker of right ventricular dysfunction in pulmonary arterial hypertension (46,47), raising the possibility that LTBP2 is a conserved mediator and indicator of fibrotic tissue remodeling across organs. Whether circulating or tissue LTBP2 predicts adverse remodeling after MI remains an important question for prospective study.

### Limitations and future directions

Our studies used global *Ltbp2*-knockout mice, which cannot exclude contributions from non-fibroblast sources of LTBP2 or developmental compensation within the microfibril system. Consistent with the latter, proteomic comparison of sham hearts showed modestly increased abundance of microfibril- and matrix-associated proteins, including MFAP4, periostin, and fibulin-1, in *Ltbp2*-null myocardium. This baseline difference was not accompanied by any detectable phenotype, sham hearts of the two genotypes were indistinguishable in interstitial collagen content, cardiomyocyte cross-sectional area, tissue stiffness, and cardiac function. Fibroblast-specific and inducible adult deletion, as has been applied to the TGF-β receptors and Smad3 (37), will nonetheless be required to establish the cell-autonomous contribution of fibroblast-derived LTBP2.

Competition between LTBP2 and LTBP1 for the fibrillin-1 N-terminus is well established biochemically (11,18,19, 36), and our findings are consistent with this mechanism operating within the assembling CFs matrix: the minimal LTBP2 module binds fibrillin-1 directly, LTBP1 fibrillar incorporation increases in the absence of LTBP2, and introducing the module acutely or by sustained expression mobilizes matrix-bound latent TGF-β. We did not, however, demonstrate that this module is itself sufficient to displace LTBP1 from fibrillin-1, nor did we assess LTBP4, which the same model predicts should be similarly affected. Both sufficiency experiments used the isolated binding module rather than full-length protein, and the recombinant and lentiviral constructs were human sequences introduced into murine fibroblasts; functional rescue indicates productive engagement with murine microfibrils despite this species difference, but direct competition assays and experiments with full-length protein will be needed to confirm displacement as the operative step.

Finally, while our in vitro data establish that LTBP2 shifts latent TGF-β from matrix-sequestered to soluble pools, the downstream activation steps remain to be defined in the intact heart. Studies of SMAD2/3 activation dynamics and the spatial distribution of latent versus active TGF-β within post-MI myocardium will be needed to connect LTBP2-dependent compartmentalization to local TGF-β activation in vivo. Our human samples were obtained from end-stage explanted hearts without regional annotation, leaving open whether LTBP2 is specifically induced in human remote myocardium after MI and whether its expression tracks with regional fibrosis or heart failure progression.

## Conclusions

This study identifies LTBP2 as a regulator of pathologic remote myocardial remodeling following myocardial infarction. LTBP2 promotes interstitial fibrosis, tissue stiffening, and functional deterioration by altering microfibrillar ECM composition and enhancing TGF-β bioavailability, while leaving the cardiomyocyte hypertrophic response intact. Genetic deletion of *Ltbp2* preserves myocardial compliance, improves regional mechanics, and enhances survival after MI without altering initial infarct size. Conversely, a minimal fibrillin-1–binding module of LTBP2 binds fibrillin-1 directly and is sufficient, acutely as purified protein and under sustained expression, to drive latent TGF-β out of the matrix and into the activatable pool. These findings establish a mechanistic link between microfibrillar organization and TGF-β–dependent profibrotic signaling, resolve how an LTBP that does not bind latent TGF-β can nonetheless amplify TGF-β signaling, and identify LTBP2 and its fibrillin-1 interaction surface as candidate therapeutic targets for preventing adverse remodeling and heart failure progression after myocardial injury.

## Author Contribution

F.E., T.N., F.S.Y., B.H., A.O.G., and S.P.H. conceived the study and designed the experiments. H.Z., M.De., M.D.P., J.L., U.K., Y.-Q.Z., T.W., H.S., A.I., J.H., C.J.R., and F.B. acquired the data or samples. F.E., M.De., M.D.P., J.L., U.K., R.H., C.J.R., and S.P.H. analyzed and interpreted the data. R.P.M. provided the *Ltbp2*-null mouse line, T.N. provided LTBP2 and FBN1 antibodies, and F.S.Y. provided transformed mink lung epithelial reporter cells and all contributed to corresponding experimental deisgns. F.E. and S.P.H. drafted the manuscript. All authors critically revised the manuscript for important intellectual content, approved the final version, and agree to be accountable for all aspects of the work.

## Novelty and Significance

### What Is Known?

- Remodeling of the remote, non-infarcted myocardium after myocardial infarction drives progressive fibrosis, ventricular stiffening, and heart failure, largely independent of infarct size.
- Transforming growth factor-β (TGF-β) is a central profibrotic signal, and its availability is controlled by latent TGF-β binding proteins (LTBPs) that tether latent complexes to fibrillin-rich microfibrils.
- LTBP2 is induced in fibrotic tissue but, unlike other family members, does not bind latent TGF-β, and how it influences TGF-β–driven fibrosis has been unresolved.

### What New Information Does This Article Contribute?

- LTBP2 is induced in injury-activated cardiac fibroblasts, deposited into fibrillin-rich matrix of both infarct and remote myocardium, and elevated in failing human left ventricle across ischemic, non-ischemic, and hypertrophic cardiomyopathy.
- Deletion of Ltbp2 improves survival, reduces remote fibrosis and tissue stiffness, and preserves regional systolic and diastolic mechanics without altering infarct size or the cardiomyocyte hypertrophic response.
- A minimal fibrillin-1–binding module of LTBP2 is sufficient to drive latent TGF-β out of the matrix and into the activatable pool, explaining how a protein that cannot bind latent TGF-β nonetheless amplifies TGF-β–driven fibrosis.

### Summary

Most therapies after myocardial infarction target the infarct itself, yet much of the decline toward heart failure originates in the surviving muscle surrounding the scar, where diffuse fibrosis stiffens the ventricle and impairs filling. We identify LTBP2 as a driver of this process. LTBP2 is produced by activated fibroblasts and incorporated into the microfibrillar scaffold of the matrix, where it reduces the amount of latent TGF-β held in reserve and increases the pool available for activation. Mice lacking LTBP2 survive infarction better and develop less fibrosis and stiffening in remote myocardium, with preserved regional mechanics but an intact hypertrophic response, indicating that the effect is specific to the interstitium. Conversely, introducing a short fibrillin-1–binding fragment of LTBP2 is enough to release latent TGF-β from the matrix within minutes. Because LTBP2 alters where TGF-β resides rather than blocking the pathway outright, and appears dispensable in the uninjured heart, targeting this interaction may offer a way to limit pathologic fibrosis without the toxicity of global TGF-β inhibition.

## Non-standard Abbreviations and Acronyms

AFM: Atomic force microscopy
APTES: 3-aminopropyltriethoxysilane
BSA: Bovine serum albumin
CF: Cardiac fibroblast
CM: Conditioned media
DIA: Data-independent acquisition
DPI: Days post-infarction
ECM: Extracellular matrix
ECV: Extracellular volume
EKV: Electrocardiogram-gated kilohertz visualization
FAC: Fractional area change
FBN1: Fibrillin-1
FS: Fractional shortening
GFP: Green fluorescent protein
GO: Gene ontology
HCM: Hypertrophic cardiomyopathy
IB: Immunoblot
ICM: Ischemic cardiomyopathy
IP: Immunoprecipitation
KO: Knockout
LA: Left atrium
LAD: Left anterior descending coronary artery
LAP: Latency-associated peptide
LLC: Large latent complex
LTBP: Latent transforming growth factor-β binding protein
LV: Left ventricle
LVEDD: Left ventricular end-diastolic diameter
LVESD: Left ventricular end-systolic diameter
MI: Myocardial infarction
NICM: Non-ischemic cardiomyopathy
OCT: Optimal cutting temperature compound
PLAX: Parasternal long-axis view
RT-qPCR: Reverse transcription quantitative polymerase chain reaction
SMAD: Mothers against decapentaplegic homolog protein
SV: Stroke volume
TCP: Tissue culture plastic
TGF-β: Transforming growth factor-β
TMLC: Transformed mink lung epithelial cell
TTC: Triphenyltetrazolium chloride
WGA: Wheat germ agglutinin
WT: Wild type

## Supplementary Figures

**Supplementary Figure 1.**
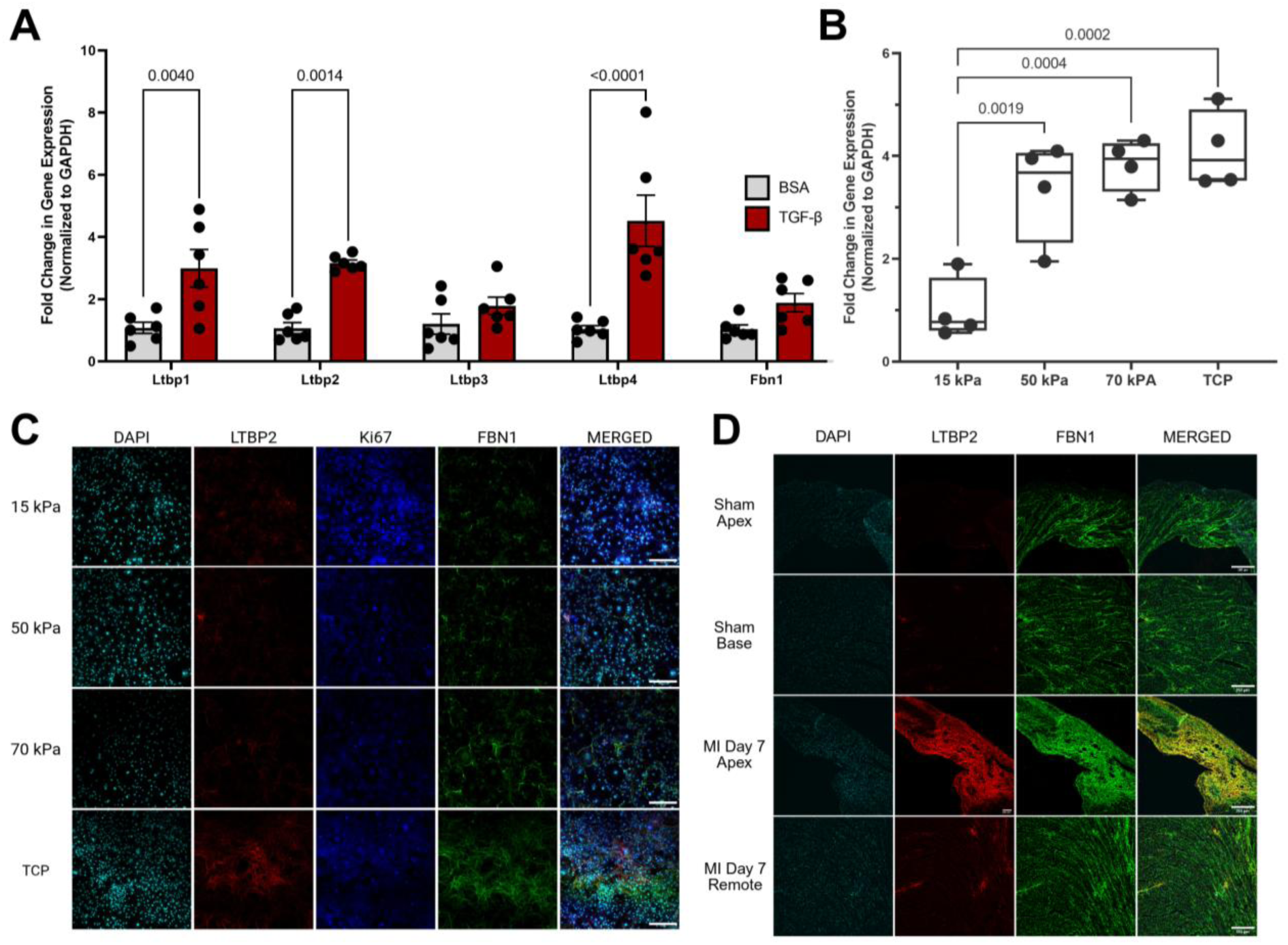
*Ltbp2* expression is induced by TGF-β stimulation and increased substrate stiffness in primary cardiac fibroblasts, and LTBP2 is deposited in infarct and remote myocardium after myocardial infarction. (A) Primary mouse cardiac fibroblasts were serum-starved for 24 hours and then stimulated with recombinant TGF-β (50 ng/mL) or bovine serum albumin (BSA; 50 ng/mL) as vehicle control for [duration]. Expression of *Ltbp1–4* and *Fbn1* was quantified by RT-qPCR, normalized to *Gapdh*, and expressed as fold change relative to BSA. n = 6 independent isolations per group. (B) *Ltbp2* expression in primary cardiac fibroblasts cultured for 7 days on silicone substrates of defined stiffness (15, 50, or 70 kPa) or on standard tissue culture plastic (TCP, >1,000 kPa), quantified by RT-qPCR and normalized to *Gapdh*. Boxes show the median and interquartile range with whiskers spanning the full range; individual replicates are overlaid. n = 4 per condition. (C) Representative immunofluorescence of primary cardiac fibroblasts cultured for 7 days under the indicated substrate stiffness conditions, stained for LTBP2 (red), Ki67 (blue), FBN1 (green), and nuclei (DAPI, cyan). LTBP2 and FBN1 deposition into fibrillar extracellular matrix structures increases with substrate stiffness. Images are representative of n = 1 independent isolations. Scale bars, 250 μm. (D) Representative immunofluorescence of mouse left ventricular myocardium stained for LTBP2 (red), FBN1 (green), and nuclei (DAPI, cyan) at 7 days after sham surgery or myocardial infarction (MI). Apical (infarct) and basal (remote) regions are shown for each condition. LTBP2 is markedly increased in the infarct and is also detectable in remote myocardium at 7 days post-MI, overlapping FBN1-positive structures. Images are representative of n = 1 hearts per group. Scale bars, [200 or 250] μm. Data in A and B are presented as indicated with individual replicates overlaid; n = 4–6 per group. Comparisons were made by two-way ANOVA with gene and treatment as factors followed by Tukey post hoc testing for **A**, and by one-way ANOVA followed by Tukey post hoc testing for **B**. All analyses were performed in GraphPad Prism.

**Supplementary Figure 2.**
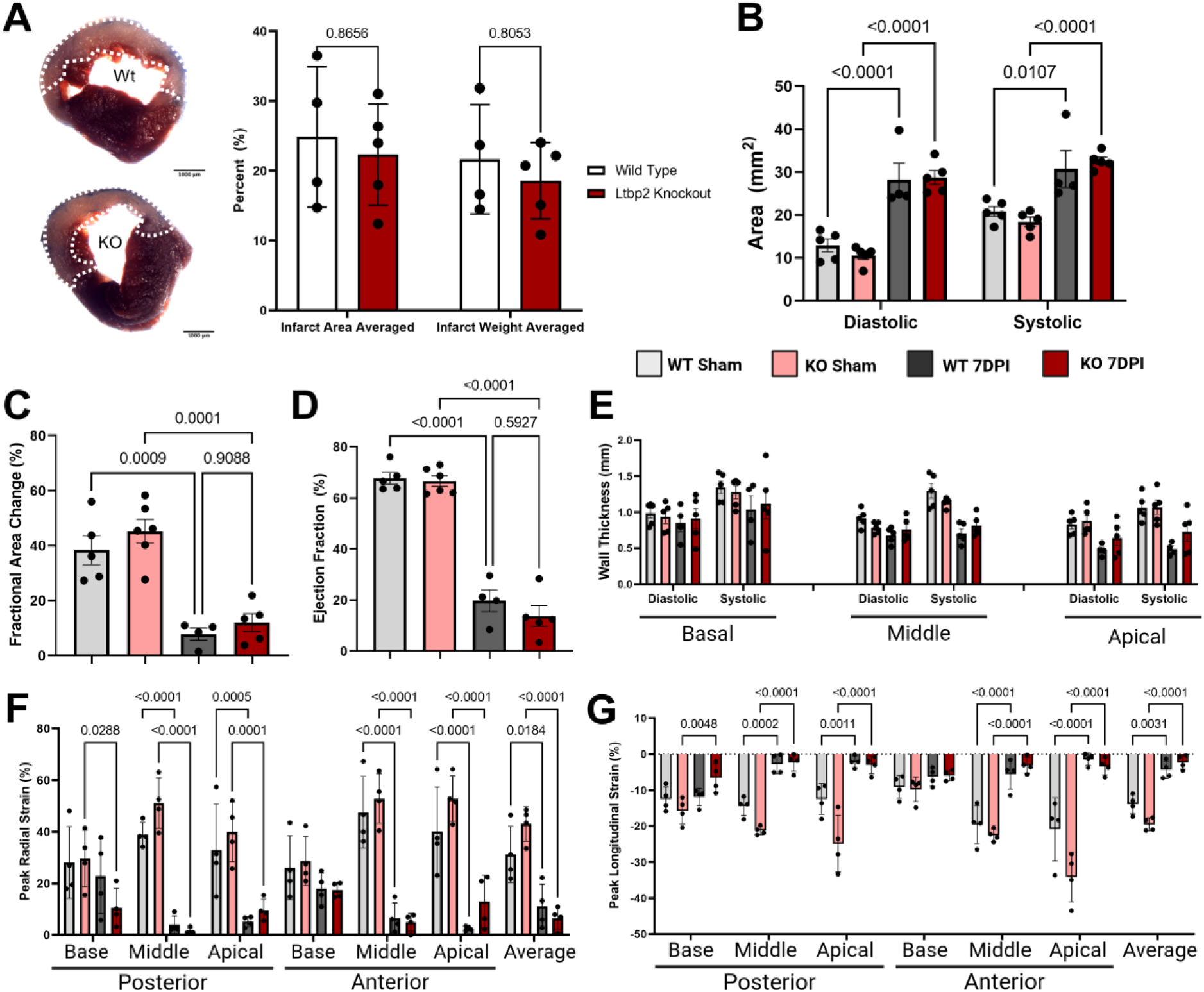
Comparable early myocardial injury and cardiac dysfunction in wild-type (WT) and *Ltbp2* knockout (KO) mice following myocardial infarction. (A) Representative triphenyltetrazolium chloride (TTC)–stained transverse heart sections from WT and KO mice at 7 days after myocardial infarction (MI), with the infarcted region outlined; viable myocardium stains red and infarcted tissue pale. Scale bars, 1,000 μm. Adjacent, Infarct area or infarct weight averaged across 5-6 sections for each heart for each genotype. (B) Two-dimensional left ventricular area at end-diastole and end-systole measured from the long-axis echocardiographic view at 7 days post-MI. (C) Fractional area change and (D) ejection fraction measured at the mid-ventricular level at 7 days post-MI. (E) Left ventricular wall thickness at end-diastole and peak systole measured at basal, mid-ventricular, and apical levels at 7 days post-MI. (F) Peak endocardial radial strain and (G) peak endocardial longitudinal strain assessed by speckle-tracking echocardiography (VevoStrain) from the long-axis view at 7 days post-MI, reported for basal, mid, and apical segments of the posterior and anterior walls and as the segment-averaged value. Groups are WT sham, KO sham, WT 7 days post-infarction (7DPI), and KO 7DPI, as indicated in the legend. Data are mean ± SEM with individual animals overlaid; n = 4-5 per group. Comparisons were made by one-way ANOVA with Tukey post hoc testing for A, C, and D, and by two-way ANOVA with group and segment (or cardiac phase) as factors followed by Tukey post hoc testing for B, E, F, and G. Only significant comparisons are annotated; no significant differences were detected between WT and KO at 7 days post-MI in any parameter. All analyses were performed in GraphPad Prism.

**Supplementary Figure 3.**
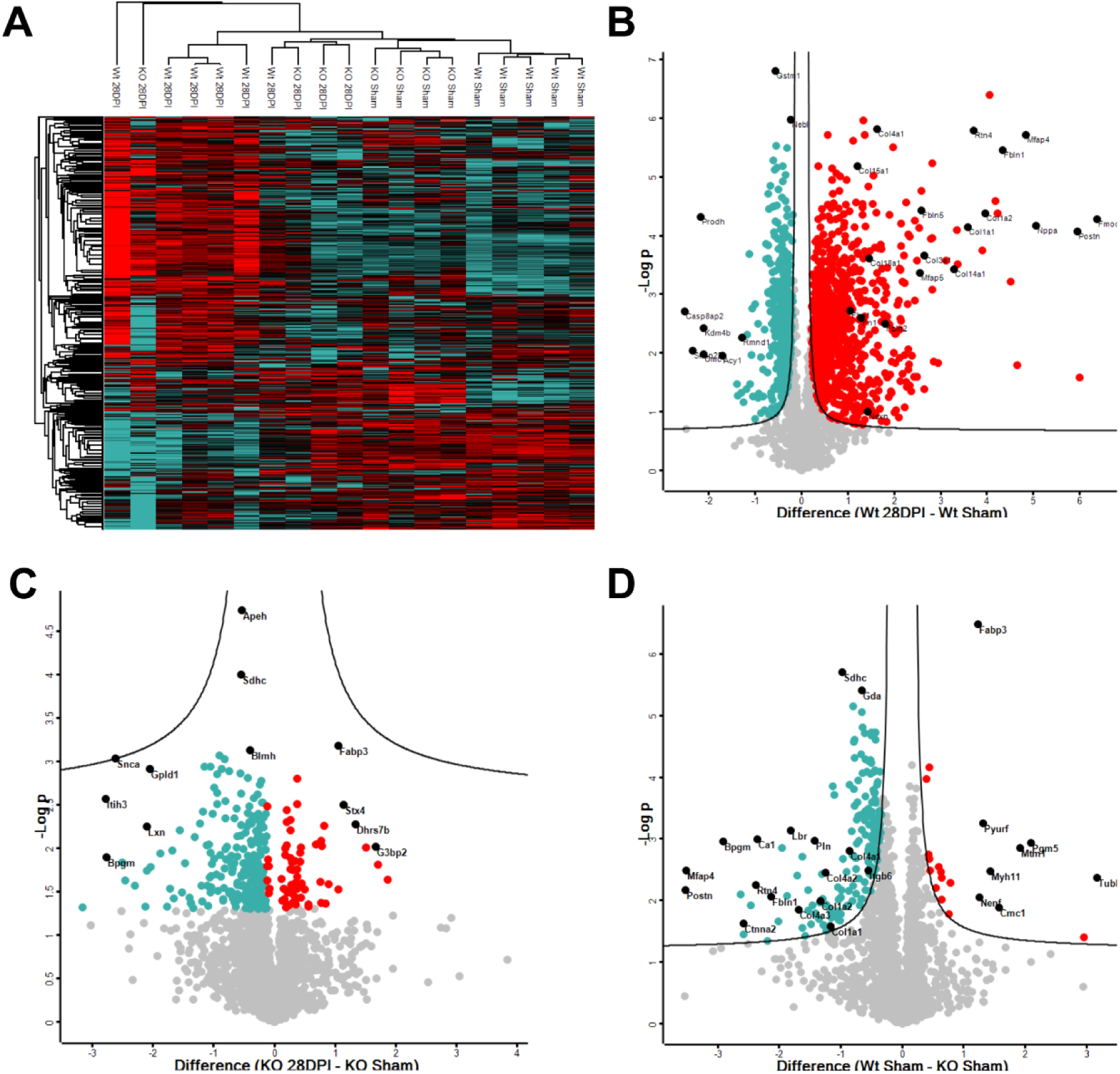
Proteomic analysis of additional group comparisons in remote myocardium of wild-type (WT) and *Ltbp2* knockout (KO) hearts at 28 days post-myocardial infarction. (A) Heat map with unsupervised hierarchical clustering of all quantified proteins across WT sham, KO sham, WT 28 days post-infarction (28DPI), and KO 28DPI remote myocardium. Rows and columns were clustered by Euclidean distance; each column represents an individual animal. Colour reflects z-scored protein intensity (teal, low; red, high). Samples segregate primarily by surgical group, with WT 28DPI forming a distinct cluster. (B) Volcano plot of differentially abundant proteins between WT 28DPI and WT sham remote myocardium, plotting the difference in mean log₂ intensity against −log₁₀ P value. Red points denote proteins significantly increased in WT 28DPI and teal points those significantly increased in WT sham. Selected extracellular matrix and fibrosis-associated proteins are labelled. (C) Volcano plot comparing KO 28DPI with KO sham remote myocardium. Since there were no significantly different proteins here based on the FDR cutoff, individual proteins are listed based on raw p-value < 0.05. (D) Volcano plot comparing WT sham with KO sham remote myocardium, presented as in B. Data are from n = 4–5 animals per group. Significance was determined by two-tailed unpaired Student’s t-test with permutation-based false discovery rate correction (FDR < 0.05, s0 = 0.01) in Perseus; curved lines indicate the significance boundary. Points falling outside the boundary are coloured; grey points are not significant

**Supplementary Figure 4.**
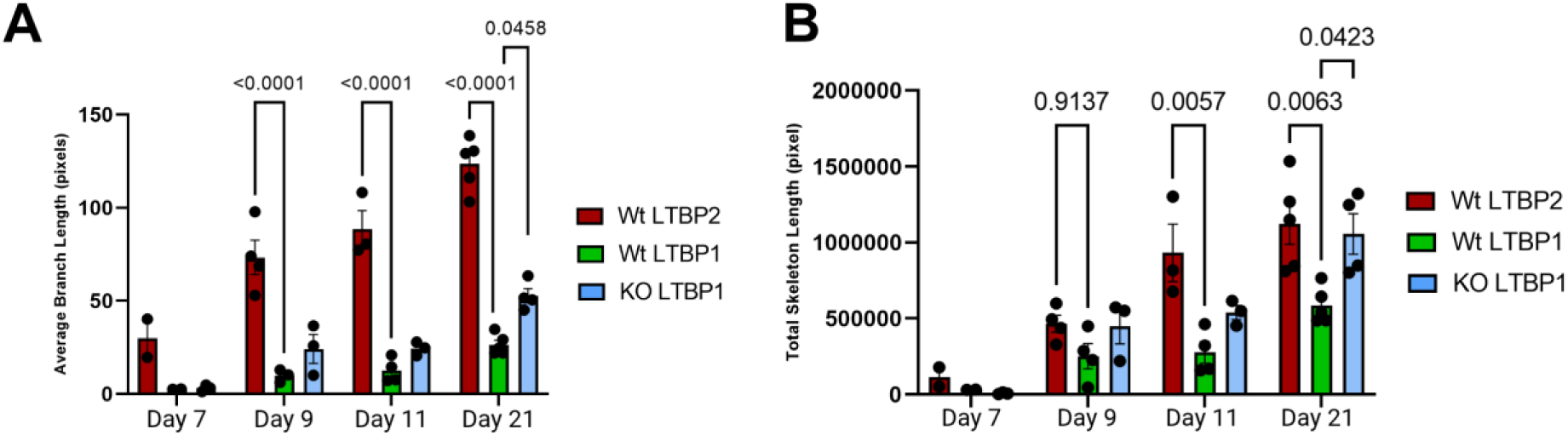
LTBP1 incorporation into fibrillar matrix networks is increased in *Ltbp2*-deficient cardiac fibroblasts. (A) Average branch length and (B) total skeleton length of LTBP1- and LTBP2-positive fibrillar structures in wild-type (WT) and *Ltbp2* knockout (KO) primary cardiac fibroblast cultures at days 7, 9, 11 and 21 after plating. Immunofluorescence images were binarized and skeletonized in ImageJ/Fiji, and branch length and total skeleton length were extracted per field as a measure of the extent and continuity of matrix deposition. Bars show WT LTBP2 (red), WT LTBP1 (green), and KO LTBP1 (blue). Data are mean ± SEM with individual replicates overlaid; n = 4-5 independent isolations per genotype and timepoint. Comparisons were made by two-way ANOVA with timepoint and stain/genotype as factors, followed by Tukey post hoc multiple comparisons testing. All analyses were performed in GraphPad Prism.

**Supplementary Figure 5.**
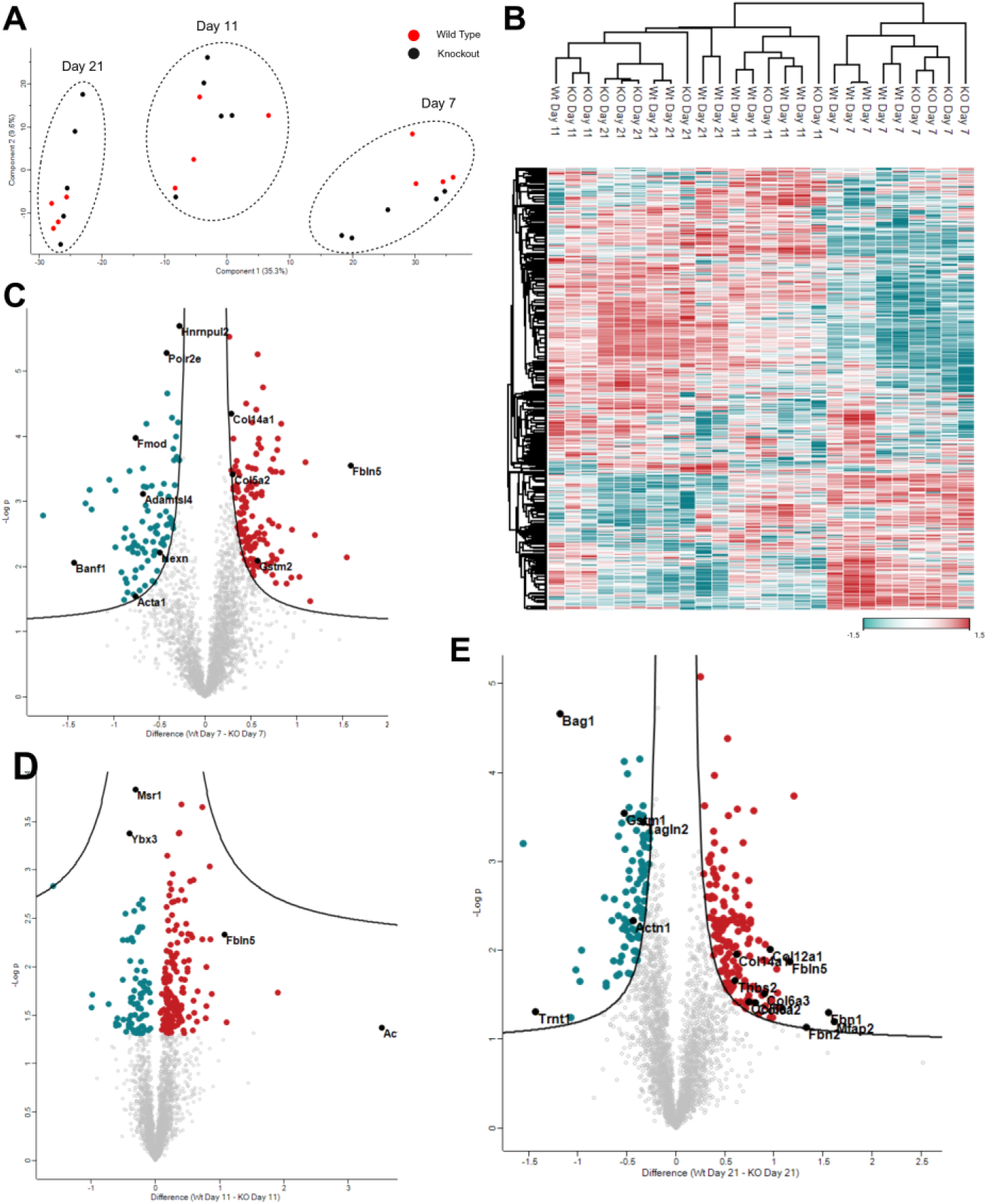
Wild-type cardiac fibroblasts deposit a more fibrotic matrix proteome than *Ltbp2*-deficient fibroblasts by 21 days in culture. (A) Principal component analysis of cell-derived extracellular matrix proteomes from wild-type (WT, red) and *Ltbp2* knockout (KO, black) primary cardiac fibroblast cultures at days 7, 11, and 21 after plating. The first two components account for 35.3% and 9.6% of total variance. Samples separate primarily by time in culture along component 1, with genotypes intermixed at each timepoint. Dashed ellipses denote timepoint groupings drawn for visualization. n = 4 for KO, n=5 for Wt independent cultures per genotype and timepoint. (B) Heat map of *Ltbp1–4* and *Fbn1* protein abundance across individual WT and KO matrix samples at days 7, 11, and 21, clustered by Euclidean distance. Colour reflects z-scored intensity across samples (green, low; red, high). (C) Volcano plot of differentially abundant matrix proteins in WT versus KO cultures at day 7, plotting the difference in mean log₂ intensity against −log₁₀ P value. Curved lines indicate the permutation-based false discovery rate boundary (FDR < 0.05, s0 = 0.1); coloured points denote proteins reaching unadjusted P < 0.05 (green, higher in WT; red, higher in KO). LTBP2 is detected in WT but not KO matrix, confirming loss of protein in the knockout. (D) Volcano plot at day 11, presented as in C. (E) Volcano plot at day 21, presented as in C, except that only proteins meeting the FDR < 0.05 threshold are coloured. Matrix and fibrosis-associated proteins including FBN1, POSTN, FMOD, THBS2, COL12A1, COL14A1, and TGFB 1 are enriched in WT matrix. Data are from n = 4 for KO, n=5 for Wts independent cultures per genotype and timepoint. Significance was determined by two-tailed unpaired Student’s t-test with permutation-based false discovery rate correction in Perseus.

**Supplementary Figure 6.**
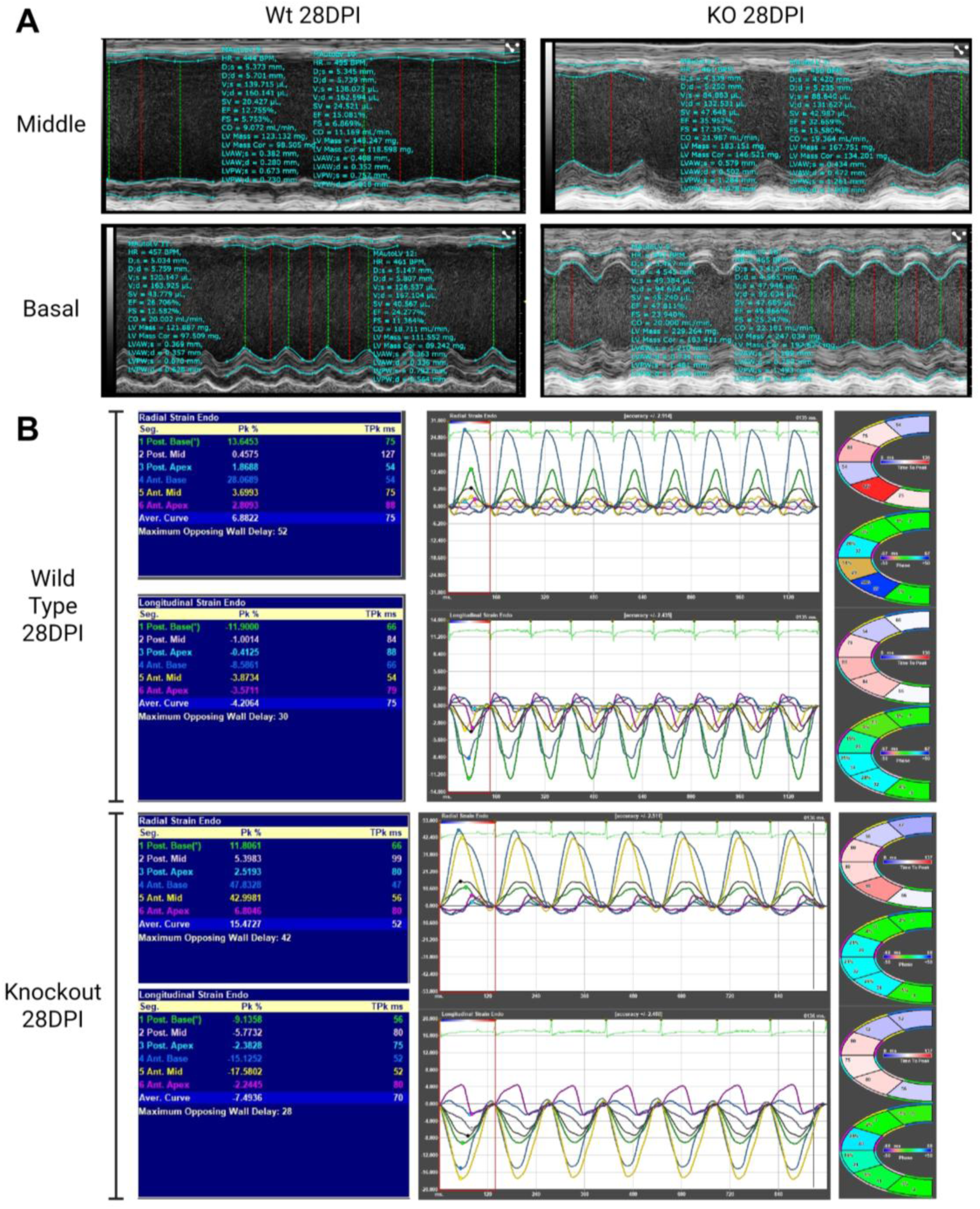
Representative M-mode and speckle-tracking strain analysis of wild-type and *Ltbp2* knockout hearts at 28 days after myocardial infarction. **(A)** Representative M-mode traces at the mid-ventricular and basal levels of the left ventricle in wild-type (WT, left) and *Ltbp2* knockout (KO, right) mice at 28 days post–myocardial infarction (MI), analyzed using the auto-LV function in VevoLab 3.1.0 with derived parameters displayed on each trace. **(B)** Representative speckle-tracking strain analysis (VevoStrain) from long-axis cines of the same hearts, showing segmental peak endocardial radial (upper) and longitudinal (lower) strain with time-to-peak values, strain-versus-time curves across consecutive cardiac cycles, and polar maps of peak strain, time to peak, and phase for the six analyzed segments. Values correspond to the parameters quantified in Figure 2D–G and representative in Tables 1-2.

## Supplementary Tables

**Table 1.** Comprehensive echocardiographic assessment of cardiac structure and function in sham and myocardial infarction groups. Echocardiographic parameters describing left ventricular (LV) morphology, systolic function, and diastolic function are presented for all experimental groups. Data are presented as mean ± SEM. Statistical comparisons between wild-type (WT) and *Ltbp2* knockout (KO) mice at 28 days post-myocardial infarction (28 dpi) were performed using two-way ANOVA followed by Tukey’s post hoc multiple comparisons test. Sample size ranged from *n* = 15–30 animals per group. No differences were detected between WT and KO sham groups for any parameters (two-way ANOVA with Tukey post-hoc, see Figure 2).

| Category of Parameter | Parameter | Wt Sham | KO Sham | Wt 28DPI | KO 28DPI | p-value | q-value | Significant? |
|---|---|---|---|---|---|---|---|---|
| 2D EKV for LV area in Long Axis view | End Systolic Area | 12.79 ± 0.69 | 12.81 ± 0.69 | 34.29 ± 2.15 | 27.65 ± 2.59 | 0.060701 | 0.085905 | No |
|  | End Diastolic Area | 22.53 ± 0.82 | 21.35 ± 0.82 | 38.43 ± 2.1 | 33.8 ± 2.42 | 0.166096 | 0.166351 | No |
|  | LV FAC | 0.43 ± 0.02 | 0.43 ± 0.02 | 0.11 ± 0.01 | 0.19 ± 0.02 | 0.003001 | 0.00888 | Yes |
| M-Mode at LV basal segment | Ejection Fraction | 61.12 ± 1.15 | 61 ± 2.7 | 33.98 ± 2.34 | 53.61 ± 3.03 | 0.000021 | 0.000277 | Yes |
|  | Fractional Wall Shortening | 32.33 ± 0.77 | 32.81 ± 2.41 | 16.43 ± 1.23 | 28.01 ± 1.97 | 0.000017 | 0.000277 | Yes |
|  | Diameter; Systole | 2.67 ± 0.09 | 2.58 ± 0.15 | 4.27 ± 0.17 | 3.37 ± 0.18 | 0.001514 | 0.005188 | Yes |
|  | Diameter; Diastole | 3.93 ± 0.1 | 3.78 ± 0.14 | 5.1 ± 0.17 | 4.67 ± 0.18 | 0.105075 | 0.121517 | No |
|  | LV Anterior Wall Thickness; Systole | 1.47 ± 0.06 | 1.55 ± 0.11 | 1.25 ± 0.1 | 1.72 ± 0.14 | 0.006813 | 0.015294 | Yes |
|  | LV Anterior Wall Thickness; Diastole | 1 ± 0.05 | 1.11 ± 0.09 | 1.01 ± 0.09 | 1.17 ± 0.08 | 0.233989 | 0.227353 | No |
|  | LV Posterior Wall Thickness; Systole | 1.24 ± 0.06 | 1.09 ± 0.05 | 1.19 ± 0.06 | 1.36 ± 0.09 | 0.123619 | 0.1364 | No |
|  | LV Posterior Wall Thickness; Diastole | 0.92 ± 0.07 | 0.84 ± 0.04 | 1.04 ± 0.05 | 1.12 ± 0.08 | 0.416685 | 0.347772 | No |
| M-Mode at LV middle segment | Ejection Fraction | 61.25 ± 1.95 | 60.85 ± 2.5 | 16.32 ± 1.33 | 35.78 ± 3.81 | 0.000007 | 0.000219 | Yes |
|  | Fractional Wall Shortening | 32.5 ± 1.52 | 32.49 ± 1.99 | 7.45 ± 0.62 | 17.52 ± 2.03 | 0.000007 | 0.000219 | Yes |
|  | Diameter; Systole | 2.69 ± 0.11 | 2.58 ± 0.11 | 5.36 ± 0.25 | 4.33 ± 0.32 | 0.016916 | 0.031464 | Yes |
|  | Diameter Diastole | 3.97 ± 0.08 | 3.8 ± 0.09 | 5.77 ± 0.24 | 5.2 ± 0.28 | 0.132959 | 0.139686 | No |
|  | LV Anterior Wall Thickness; Systole | 1.34 ± 0.06 | 1.33 ± 0.08 | 0.6 ± 0.05 | 1.03 ± 0.11 | 0.000564 | 0.002822 | Yes |
|  | LV Anterior Wall Thickness; Diastole | 0.91 ± 0.04 | 1.53 ± 0.62 | 0.51 ± 0.04 | 0.76 ± 0.09 | 0.00915 | 0.018614 | Yes |
|  | LV Posterior Wall Thickness; Systole | 1.18 ± 0.03 | 1.15 ± 0.07 | 0.9 ± 0.08 | 1.14 ± 0.08 | 0.058232 | 0.085905 | No |
|  | LV Posterior Wall Thickness; Diastole | 0.82 ± 0.03 | 0.81 ± 0.03 | 0.78 ± 0.07 | 0.89 ± 0.07 | 0.318282 | 0.283193 | No |
| M-Mode in short axis view at LV middle segment | Ejection Fraction | 58.56 ± 1.95 | 60.65 ± 1.62 | 18.21 ± 1.25 | 30.22 ± 3.06 | 0.00023 | 0.001665 | Yes |
|  | Fractional Wall Shortening | 30.79 ± 1.37 | 31.6 ± 1.17 | 8.39 ± 0.63 | 14.44 ± 1.55 | 0.000275 | 0.00179 | Yes |
| B-mode EKV in short Axis Area at LV middle segment | End Systolic Area | 5.46 ± 0.49 | 5.06 ± 0.48 | 19.16 ± 1.32 | 15.32 ± 2.42 | 0.140984 | 0.144294 | No |
|  | End Diastolic Area | 12.66 ± 0.59 | 12.04 ± 0.77 | 24.76 ± 1.17 | 23.26 ± 1.81 | 0.472092 | 0.379422 | No |
|  | LV FAC | 0.58 ± 0.03 | 0.58 ± 0.02 | 0.23 ± 0.03 | 0.37 ± 0.06 | 0.022663 | 0.039002 | Yes |
| M-Mode in short axis view at LV middle segment | LV Anterior Wall Thickness; Systole | 1.22 ± 0.06 | 1.28 ± 0.06 | 0.72 ± 0.07 | 0.82 ± 0.08 | 0.416375 | 0.347772 | No |
|  | LV Anterior Wall Thickness; Diastole | 0.85 ± 0.05 | 0.89 ± 0.05 | 0.64 ± 0.07 | 0.65 ± 0.09 | 0.937465 | 0.642841 | No |
|  | LV Posterior Wall Thickness; Systole | 1.14 ± 0.05 | 1.19 ± 0.06 | 0.98 ± 0.06 | 1.12 ± 0.13 | 0.249221 | 0.238592 | No |
|  | LV Posterior Wall Thickness; Diastole | 0.76 ± 0.03 | 0.85 ± 0.04 | 0.85 ± 0.06 | 0.89 ± 0.1 | 0.669524 | 0.512777 | No |
|  | Diameter; Systole | 2.8 ± 0.1 | 2.64 ± 0.08 | 5.07 ± 0.17 | 4.68 ± 0.36 | 0.271119 | 0.252141 | No |
|  | Diameter Diastole | 4.03 ± 0.08 | 3.85 ± 0.07 | 5.55 ± 0.16 | 5.44 ± 0.35 | 0.749358 | 0.554355 | No |
| Mitral Valve Hemodynamics | E-Wave | 628.06 ± 19.46 | 666.08 ± 19.7 | 556.7 ± 22.01 | 655.3 ± 51.82 | 0.0492 | 0.07626 | No |
|  | A-Wave | 449.03 ± 11.33 | 491.22 ± 15.53 | 367.26 ± 40.25 | 379.41 ± 34.92 | 0.847355 | 0.606185 | No |
|  | e' Wave | 24.76 ± 1.02 | 30.03 ± 1.42 | 22.31 ± 1.57 | 24.3 ± 2.96 | 0.517691 | 0.410996 | No |
|  | E/A Ratio | 1.41 ± 0.05 | 1.37 ± 0.06 | 1.9 ± 0.28 | 1.9 ± 0.31 | 0.992215 | 0.665909 | No |
|  | E/e' Ratio | 25.68 ± 0.82 | 22.61 ± 0.98 | 25.81 ± 1.52 | 29.4 ± 3.24 | 0.262608 | 0.247765 | No |
|  | Total Systolic Time (IVCT+ET+IVRT) | 77.8 ± 2.23 | 81.11 ± 1.97 | 82.76 ± 2.18 | 84 ± 1.15 | 0.70552 | 0.527923 | No |
|  | Ejection Time | 53.83 ± 1.23 | 56.73 ± 1.95 | 48.28 ± 1.59 | 55.05 ± 1.27 | 0.01088 | 0.021463 | Yes |
|  | MPI (IVCT+IVRT/ET) | 0.45 ± 0.03 | 0.44 ± 0.02 | 0.73 ± 0.04 | 0.53 ± 0.03 | 0.002704 | 0.008383 | Yes |

**Table 2.** Comprehensive echocardiographic speckle tracking strain assessment of cardiac structure and function in sham and myocardial infarction groups. Echocardiographic parameters describing left ventricular (LV) strain presented in systolic function, and diastolic function are presented for all experimental groups. Data are presented as mean ± SEM. Statistical comparisons between wild-type (WT) and *Ltbp2* knockout (KO) mice at 28 days post-myocardial infarction (28 dpi) were performed using two-way ANOVA followed by Tukey’s post hoc multiple comparisons test. Sample size ranged from *n* = 15–30 animals per group. No differences were detected between WT and KO sham groups for any parameters (two-way ANOVA with Tukey post-hoc, see Figure 2).

| Category of Parameter | Parameter | Wt Sham | KO Sham | Wt 28DPI | KO 28DPI | p-value | q-value | Significant? |
|---|---|---|---|---|---|---|---|---|
| Radial Strain in long axis view of LV | Posterior Basal Wall | 31.46 ± 1.34 | 21.24 ± 1.3 | 15.31 ± 2.45 | 25.53 ± 3.35 | 0.019758 | 0.035729 | Yes |
|  | Posterior Middle Wall | 44.68 ± 2.46 | 31.7 ± 3 | 6.4 ± 1.38 | 13.12 ± 3.74 | 0.077165 | 0.106882 | No |
|  | Posterior Apical Wall | 30.75 ± 4.3 | 25.27 ± 3.01 | 6.19 ± 1.69 | 7.33 ± 2.24 | 0.682053 | 0.516298 | No |
|  | Anterior Basal Wall | 25.3 ± 3.02 | 20.19 ± 1.39 | 12.29 ± 2.25 | 24.03 ± 2.02 | 0.001151 | 0.004163 | Yes |
|  | Anterior Middle Wall | 39.74 ± 2.23 | 29.66 ± 3.05 | 8.52 ± 2.65 | 21.62 ± 3.79 | 0.008132 | 0.017077 | Yes |
|  | Anterior Apical Wall | 27.15 ± 4.22 | 23 ± 2.78 | 6.52 ± 1.48 | 7.91 ± 2.08 | 0.581421 | 0.45603 | No |
|  | Average Strain | 31.01 ± 2.22 | 23.97 ± 1.91 | 7.51 ± 1.53 | 15.01 ± 1.89 | 0.005157 | 0.013429 | Yes |
| Longitudinal Strain in long axis view of LV (-) | Posterior Basal Wall | 14.75 ± 1.44 | 10.55 ± 1.27 | 7.36 ± 0.82 | 12.51 ± 0.6 | 0.000096 | 0.000894 | Yes |
|  | Posterior Middle Wall | 15.78 ± 1.26 | 14.52 ± 1.01 | 4.03 ± 1 | 7.3 ± 1.19 | 0.046748 | 0.074226 | No |
|  | Posterior Apical Wall | 13.81 ± 2.31 | 16.63 ± 1.49 | 2.49 ± 0.73 | 4.97 ± 1.38 | 0.106091 | 0.121517 | No |
|  | Anterior Basal Wall | 12.37 ± 0.79 | 7.7 ± 1.06 | 6.96 ± 0.92 | 8.67 ± 1.35 | 0.28927 | 0.265232 | No |
|  | Anterior Middle Wall | 21.96 ± 2.57 | 18.95 ± 1.68 | 4.52 ± 1.17 | 11.15 ± 1.94 | 0.005703 | 0.013896 | Yes |
|  | Anterior Apical Wall | 19.06 ± 2.13 | 24.21 ± 1.72 | 4.91 ± 1.57 | 7.47 ± 2.03 | 0.32191 | 0.283193 | No |
|  | Average Strain | 14.06 ± 1.62 | 13.96 ± 0.87 | 4.1 ± 0.58 | 8.2 ± 0.8 | 0.000358 | 0.002117 | Yes |
| Systolic Radial Strain Rate in long axis view of LV | Posterior Basal Wall | 7.42 ± 0.44 | 4.85 ± 0.59 | 4.03 ± 0.54 | 6.04 ± 0.66 | 0.025829 | 0.042037 | Yes |
|  | Posterior Middle Wall | 9.57 ± 0.45 | 7.55 ± 0.69 | 2.23 ± 0.3 | 3.44 ± 0.79 | 0.133034 | 0.139686 | No |
|  | Posterior Apical Wall | 7.27 ± 0.89 | 6.4 ± 0.67 | 2.43 ± 0.47 | 2.62 ± 0.47 | 0.786326 | 0.575166 | No |
|  | Anterior Basal Wall | 5.44 ± 0.57 | 4.79 ± 0.41 | 4.27 ± 0.53 | 5.53 ± 0.42 | 0.092322 | 0.120374 | No |
|  | Anterior Middle Wall | 9.2 ± 0.53 | 7.28 ± 0.82 | 3.13 ± 0.53 | 5.13 ± 0.87 | 0.051954 | 0.078657 | No |
|  | Anterior Apical Wall | 7 ± 0.51 | 6.34 ± 0.59 | 2.49 ± 0.39 | 2.96 ± 0.44 | 0.434645 | 0.353693 | No |
|  | Average Strain | 7.1 ± 0.46 | 5.95 ± 0.43 | 1.99 ± 0.38 | 4.48 ± 0.53 | 0.000753 | 0.003267 | Yes |
| Diastolic Radial Strain Rate in long axis view of LV (-) | Posterior Basal Wall | 6.82 ± 0.63 | 5.94 ± 0.65 | 4.51 ± 0.5 | 6.89 ± 0.73 | 0.011464 | 0.021951 | Yes |
|  | Posterior Middle Wall | 8.81 ± 0.69 | 7.4 ± 0.69 | 2.15 ± 0.31 | 4.5 ± 0.74 | 0.004524 | 0.01227 | Yes |
|  | Posterior Apical Wall | 6.71 ± 1.26 | 6.46 ± 0.74 | 2.69 ± 0.79 | 2.9 ± 0.47 | 0.831041 | 0.60112 | No |
|  | Anterior Basal Wall | 6.14 ± 0.74 | 6.72 ± 0.58 | 3.6 ± 0.43 | 6.25 ± 0.75 | 0.00407 | 0.011521 | Yes |
|  | Anterior Middle Wall | 8.59 ± 0.98 | 7.34 ± 0.68 | 2.25 ± 0.39 | 5.47 ± 0.82 | 0.000994 | 0.003805 | Yes |
|  | Anterior Apical Wall | 6.33 ± 1.22 | 5.67 ± 0.7 | 2.76 ± 0.41 | 3.05 ± 0.36 | 0.612847 | 0.474956 | No |
|  | Average Strain | 6.94 ± 0.83 | 6.1 ± 0.55 | 1.73 ± 0.27 | 4.41 ± 0.7 | 0.000716 | 0.003267 | Yes |
| Systolic Longitudinal Strain Rate in long axis view of LV (-) | Posterior Basal Wall | 4.52 ± 0.42 | 3.01 ± 0.53 | 2.17 ± 0.25 | 3.3 ± 0.3 | 0.007865 | 0.017068 | Yes |
|  | Posterior Middle Wall | 5.01 ± 0.37 | 4.31 ± 0.35 | 1.4 ± 0.37 | 2.32 ± 0.32 | 0.082015 | 0.111233 | No |
|  | Posterior Apical Wall | 4.33 ± 0.79 | 4.82 ± 0.81 | 0.99 ± 0.18 | 1.74 ± 0.36 | 0.059538 | 0.085905 | No |
|  | Anterior Basal Wall | 3.6 ± 0.74 | 3.81 ± 0.95 | 2.59 ± 0.37 | 2.67 ± 0.42 | 0.883733 | 0.625337 | No |
|  | Anterior Middle Wall | 6.83 ± 0.59 | 5.9 ± 0.76 | 1.51 ± 0.33 | 4.36 ± 0.72 | 0.000885 | 0.003599 | Yes |
|  | Anterior Apical Wall | 6.19 ± 0.7 | 7.94 ± 0.71 | 1.94 ± 0.47 | 3.29 ± 0.64 | 0.094302 | 0.120374 | No |
|  | Average Strain | 4.66 ± 0.48 | 4.46 ± 0.41 | 1.42 ± 0.24 | 2.04 ± 0.32 | 0.130993 | 0.139686 | No |
|  | Maximum Opposing Wall Delay | 40.71 ± 14.16 | 34.57 ± 10.36 | 42.5 ± 5.11 | 34.7 ± 8.61 | 0.427498 | 0.35228 | No |
| Diastolic Longitudinal Strain Rate in long axis view of LV | Posterior Basal Wall | 4.57 ± 0.37 | 4.73 ± 0.36 | 2.07 ± 0.24 | 4 ± 0.26 | 0.000021 | 0.000277 | Yes |
|  | Posterior Middle Wall | 4.67 ± 0.55 | 4.96 ± 0.52 | 1.26 ± 0.26 | 2.56 ± 0.34 | 0.005763 | 0.013896 | Yes |
|  | Posterior Apical Wall | 4.02 ± 0.76 | 5.35 ± 0.7 | 0.96 ± 0.19 | 2.01 ± 0.42 | 0.022766 | 0.039002 | Yes |
|  | Anterior Basal Wall | 4.72 ± 0.59 | 4.95 ± 0.66 | 2.23 ± 0.27 | 3.76 ± 0.25 | 0.000549 | 0.002822 | Yes |
|  | Anterior Middle Wall | 0.06 ± 0.01 | 0.06 ± 0 | 0.01 ± 0 | 0.04 ± 0.01 | 0.00015 | 0.001218 | Yes |
|  | Anterior Apical Wall | 0.06 ± 0.01 | 0.09 ± 0.01 | 0.02 ± 0 | 0.03 ± 0.01 | 0.104842 | 0.121517 | No |
|  | Average Strain | 0.05 ± 0.01 | 0.05 ± 0 | 0.01 ± 0 | 0.03 ± 0 | 0.000056 | 0.000608 | Yes |
|  | Maximum Opposing Wall Delay | 26 ± 11.17 | 18.63 ± 6.79 | 70.92 ± 9.55 | 45.6 ± 10.83 | 0.094218 | 0.120374 | No |

